# The oligomannose N-glycans 3D architecture and its response to the FcγRIIIa structural landscape

**DOI:** 10.1101/2021.01.11.426234

**Authors:** Carl A Fogarty, Elisa Fadda

## Abstract

Oligomannoses are evolutionarily the oldest class of N-glycans, where the arms of the common pentasaccharide unit, i.e. Man α1-6)-[Manα(1-3)]-Manβ(1-4)-GlcNAcβ(1-4)-GlcNAcβ1-Asn, are functionalized exclusively with branched arrangements of mannose (Man) monosaccharide units. In mammalian species oligomannose N-glycans can have up to 9 Man, meanwhile structures can grow to over 200 units in yeast mannan. The highly dynamic nature, branching complexity and 3D structure of oligomannoses have been recently highlighted for their roles in immune escape and infectivity of enveloped viruses, such as HIV-1 and SARS-CoV2. The architectural features that allow these N-glycans to perform their functions is yet unclear, due to their intrinsically disordered nature that hinders their structural characterization. In this work we will discuss the results of over 54 µs of cumulative sampling by molecular dynamics (MD) simulations of differently processed, free (not protein-linked) oligomannose N-glycans common in vertebrates. We then discuss the effects of a protein surface on their structural equilibria based on over 4 µs cumulative MD sampling of the fully glycosylated CD16a Fc gamma receptor (FcγRIIIa), where the type of glycosylation is known to modulate its binding affinity for IgG1s, regulating the antibody-dependent cellular cytotoxicity (ADCC). Our results show that the protein’s structural constraints shift the oligomannoses conformational ensemble to promote conformers that satisfy the steric requirements and hydrogen bonding networks demanded by the protein’s surface landscape. More importantly, we find that the protein does not actively distort the N-glycans into structures not populated in the unlinked forms in solution. Ultimately, the highly populated conformations of the Man5 linked glycans support experimental evidence of high levels of hybrid complex forms at N45 and show a specific presentation of the arms at N162, which may be involved in mediating binding affinity to the IgG1 Fc.

## Introduction

Complex carbohydrates (or glycans) are the most abundant biomolecules in nature. Within a human biology context, glycans coat cell membranes and protein surfaces, mediating a myriad of essential biological processes in health and disease states^1-6^. N-glycosylation is one of the most abundant and diverse type of post-translational modification that can affect protein trafficking, structural stability and mediate interactions with different receptors^6-11^. N-glycan recognition and binding affinities are often highly specific to their sequence, intended as the types of monosaccharides, their stereochemistry and branching patterns^12^; a principle that has been successfully exploited in the development of glycan microarray technology^13^.

Molecular recognition is fundamentally dependent, among other considerations, on structural and electrostatic complementarity between the ligand and the receptor’s binding site. Within this framework, the prediction and characterization of glycan binding specificity is an extremely difficult task, due to their high degree of flexibility or intrinsic disorder, which hinders our ability to determine their 3D structure by means of experimental techniques. Indeed, glycans can only be structurally resolved in their entirety only when tightly bound to a receptor, thus when their conformational degrees of freedom are heavily restrained. Because of their inherent flexibility, free glycans can adopt different 3D structures within a weighted conformational ensemble, which cannot be determined with currently available experimental methods; although very promising steps forward have been recently made in advancing imaging techniques for single glycans^14, 15^.

High performance computing (HPC) molecular simulations can contribute a great deal towards our understanding of the relationships between glycans’ sequence, structure and function. Indeed, conformational sampling through conventional and/or enhanced molecular dynamics (MD) schemes allows us to characterize the dynamic behaviour of different glycoforms at the atomistic level of details. Within this context, for the past few years our lab contributed to the knowledge of N-glycans dynamics by providing information on their 3D architecture and relative flexibility from extensive MD-based conformational sampling^16, 17^. As an example, we have shown how the sequence (and branching) of complex N-glycans determines the 3D structure, which in turn drives their recognition^16, 17^. In this work we extend our dataset of free (unlinked) N-glycans structures to the vertebrate oligomannose type, where, as shown in **Figure 1**, the common pentasaccharide unit, i.e. Manα (1-6)-[Manα (1-3)]-Manβ(1-4)-GlcNAcβ(1-4)-GlcNAcβ1-Asn, is functionalised by a branched arrangement of only Man units. In addition, we also determine how the protein surface landscape affects their conformational dynamics, which is a very important question in terms of its impact on molecular recognition and function, while challenging to answer in absolute terms because of the site-specific character.

**Figure 1.**
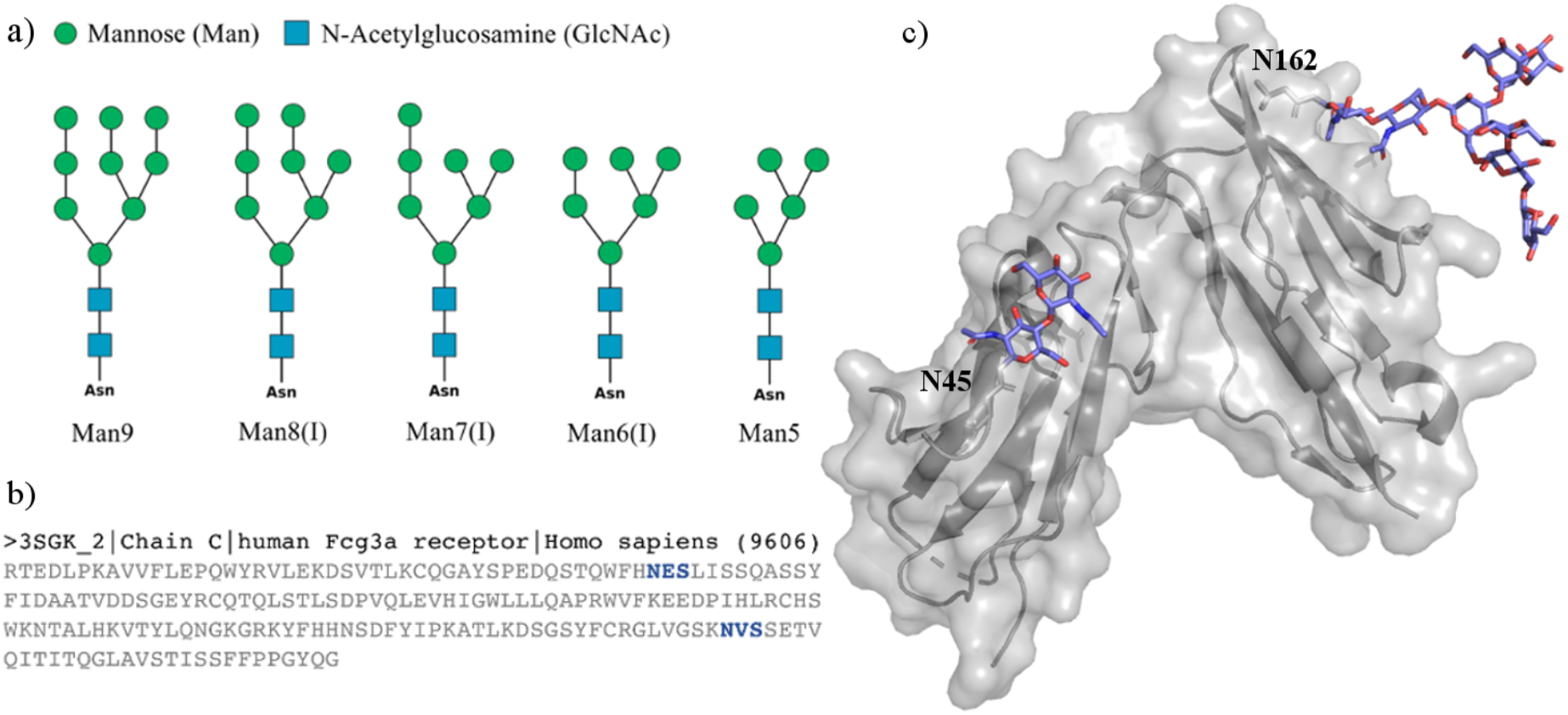
**Panel a)** SNFG representation^18^ of a subset of the oligomannose N-glycans discussed in this work. Man6/7/8(I) indicate specific positional isomers; for the complete list of isomers see Figure S.1. **Panel b)** Sequence of the human CD16a (FcγIIIa) of the PDB 3SGK^19^ with the occupied sequons highlighted in blue. **Panel c)** Structure of the human CD16a (FcγIIIa) from PDB 3SGK with the resolved N-glycans at positions N45 and N162 represented with sticks. Image rendered with VMD (www.vmd.org). N-glycan sketches rendered with DrawGlycan (http://www.virtualglycome.org/DrawGlycan/).

Oligomannoses are often defined as “immature” N-glycans, as they are processed towards complex functionalization in the Golgi^6^ and are not abundant in vertebrates. Nevertheless, these N-glycans are a common post-translational modification of viral envelope proteins expressed in human cell lines^20, 21^, for example it is the prevalent type of glycosylation of the HIV-1 fusion trimer^22-25^. Furthermore, an increase in large oligomannose-type N-glycosylation in humans has been linked to breast cancer progression^26-28^ and can occur where the protein landscape at the N-glycan site does not allow easy access to the required glycohydrolases and glycotransferases for further functionalization^6, 29, 30^. Interestingly, recent work has shown that oligomannose N-glycans functionalizing CD16a low-affinity Fc γ receptors (FcγIIIa) determine an increase in IgG1-binding affinity by 51-fold^31^, relative to the more common complex N-glycans^32^, although the N-glycosylation composition varies depending on the glycosylation site^32^.

In this work we have studied the effect of the FcγIIIa protein surface landscape on the intrinsic conformational propensity of different oligomannose N-glycans we determined for the unlinked forms. Our results show that the two FcγIIIa N-glycosylation sites, N45 and N162, affect the oligomannose dynamics rather differently, in function of the structural constrains of the sites and of the 3D architecture of the glycan. More specifically, we find that the protein landscape affects the glycans conformational equilibrium by promoting structures that are complementary to it and not by actively changing their intrinsic architecture. Indeed, all the 3D conformers observed in the analysis of the bound oligomannoses, are always identified in the simulations of the corresponding unlinked forms in solution, although in different populations. Interestingly, we also determined that the progressive elongation of the arms/branches promotes inter-arm contacts, where the Man9 3D architecture is almost entirely structured with interacting arms. Finally, these findings fit very well within the framework of our recently proposed “glycoblocks” glycans structure representation^16^, whereby groups of specifically linked monosaccharides within N-glycans represent independent structural elements (or glycoblocks), which exposure, or presentation in function of the particular protein landscape, drives molecular recognition.

### Computational Method

All 12 oligomannose starting structures, shown in **Figures 1** and **S.1**, were obtained with the GLYCAM carbohydrate builder online tool (http://www.glycam.org). For each of these oligomannoses, we built nine structures characterized by different combinations of the two α (1-6) torsions values. Complete topology and parameter files where generated with the *tLEaP* tool from version 18 of the AMBER software package^33^, with the GLYCAM06-j1 parameter set^34^ to represent the carbohydrates and the TIP3P model^35^ for water molecules. Because our simulations do not involve the calculation of hydration or of carbohydrate-carbohydrate binding free energies^36, 37^, and also because of consistency with our previous work^16, 17^, we consider the choice of GLYCAM06-j1/TIP3P parameter set as appropriate. All simulations were run in 200 mM NaCl salt concentration, with counterions represented by amber parameters^38^ in a cubic simulation box of 16 Å sides. Long range electrostatic were treated by Particle Mesh Ewald (PME) with cut-off set at 11 Å and a B-spline interpolation for mapping particles to and from the mesh of order of 4. Van der Waals (vdW) interactions were cut-off at 11 Å. The MD trajectories were generated by Langevin dynamics with collision frequency of 1.0 ps^-1^. Pressure was kept constant by isotropic pressure scaling with a pressure relaxation time of 2.0 ps. After an initial 500.000 cycles of steepest descent energy minimization, with all protein/glycans heavy atoms restrained by a harmonic potential with a force constant of 5 kcal mol^-1^Å^-2^, the system was heated in two stages, i.e. from 0 to 100 K over 500 ps at constant volume and then from 100 K to 300 K over 500 ps at constant pressure. After the heating phase all restraints were removed and the system was allowed to equilibrate for 5 ns at 300 K and at 1 atm of pressure. Production and subsequent analysis was done on 500 ns trajectories run in parallel for each uncorrelated starting structure, i.e. each conformer generated with GLYCAM-Web. Analysis was done using the *cpptraj* tool and with VMD^39^ (https://www.ks.uiuc.edu/Research/vmd/). The dihedral distributions from the trajectories were obtained in terms of kernel density estimates (KDE), with a smoothing parameter of [1000,1000], with the ks package in R and rendered with heat maps with RStudio (www.rstudio.com) in conjunction with the DBSCAN clustering algorithm. The highest populated conformers resulting from the analysis of Man5 and Man9 were then grafted in positions N45 and N162 of the FcgRIIIa (PDB 3SGK) by structural alignment to the resolved chitobiose at N45, see **Figure 1c**, and to the N162 sidechain, to obtain two systems, one with only Man5 and the other with only Man9 at both positions. As a note, the structure of the N-glycan at N165 from the crystal structure is quite distorted with uncommon conformations of some of the monosaccharides, probably determined by the fitting to the electron density, therefore it was disregarded and only the chitobiose was used for structural alignment. These systems were run in duplicates from uncorrelated starting structures with the same simulation protocol used for the free glycans. Production runs were extended to 1 μs for each trajectory, for a total of 4 μs of cumulative sampling time. All simulations were run on NVIDIA Tesla V100 16GB PCIe (Volta architecture) GPUs on resources from the Irish Centre for High-End Computing (ICHEC) (www.ichec.ie).

## Results

We used conventional MD simulations, run in parallel for 500 ns from nine uncorrelated starting points^16, 17^, to characterize the 3D structure and dynamics of human oligomannose N-glycans, when unlinked, see **Figures 1** and **S.1**. The effects of the protein on their intrinsic dynamics was studied on two models with Man5 and Man9 linked to the human FcγIIIa protein on the two N-glycosylation sites, namely N45 and N162, see **Figure 1c**. This section is organized as follows, first we present the results obtained for the unlinked oligomannoses, starting with Man5 that we used as a reference to describe sequence-to-structure changes in the larger forms. The subset of representative isomers shown in **Figure 1** is presented here for simplicity, while the complete analysis of all positional isomers with heat maps and tables is included as Supplementary Material. The section concludes with the results obtained for Man5 and Man9 when linked to the FcγRIIIa. **Man5** is the simplest oligomannose found in vertebrates and the substrate of GlcNAc transferase I (GnTI), responsible to start the N-glycan complex functionalization in the Golgi^6^. As found for complex biantennary N-glycans^16, 17^ the Man5 chitobiose core and the following Manβ(1-4)-GlcNAc linkage are rigid with only one conformation significantly occupied, see **Figure 2** and **Table S.1**, while the (1-3) arm adopts an outstretched conformation with flexibility in a range of 40° around the psi torsion angle, see **Figure 2** and **Table 1**. The Man5 (1-6) arm has a relatively more complex dynamics, hinging around the preferential ‘open’ conformation^16, 17^, populated at 82%, where the Manα (1-3)-Man branch can be orientated towards the front of the page and the Manα (1-6)-Man branch towards the back of the page, or *vice versa*. We also identified two alternative, less populated conformers, namely a “front fold” (phi = 79°, psi = 87°) with a relative population of 12% and a “back fold” (phi = 83°, psi = - 76°) with a relative population of 6%, see **Figure 2** and **Table 1**. In the front fold the terminal Manα (1-6)-Man interacts through hydrogen bonds with the N-acetyl group of the second core GlcNAc, pushing Manα (1-3)-Man upwards, while the back fold is stabilized by hydrogen bonds between the terminal Manα (1-6)-Man and both GlcNAc residues of the chitobiose, see **Figure 2.b**. The open conformation can be further analysed in terms of the α (1-6) linkage omega torsion angle that determines different orientations of the Manα (1-6)-Man and Manα (1-3)-Man branches relative to the core. As shown in **Figure 3**, there are two dominant conformers contributing to the open structure, open (cluster 1) populated at 48% with omega = 56° and open (cluster 2) populated at 33% with omega = -175°, see **Table 1**. The dynamics of the branches on the Man5 (1-6) arm follows the same pattern observed in the (1-3) and (1-6) arms, with only small differences dictated by their immediate environment. For example, the terminal Manα (1-6)-Man is found predominantly (80%) in the open (cluster 1) conformation, see **Table 1**, and does not show a back fold orientation.

**Table 1.**
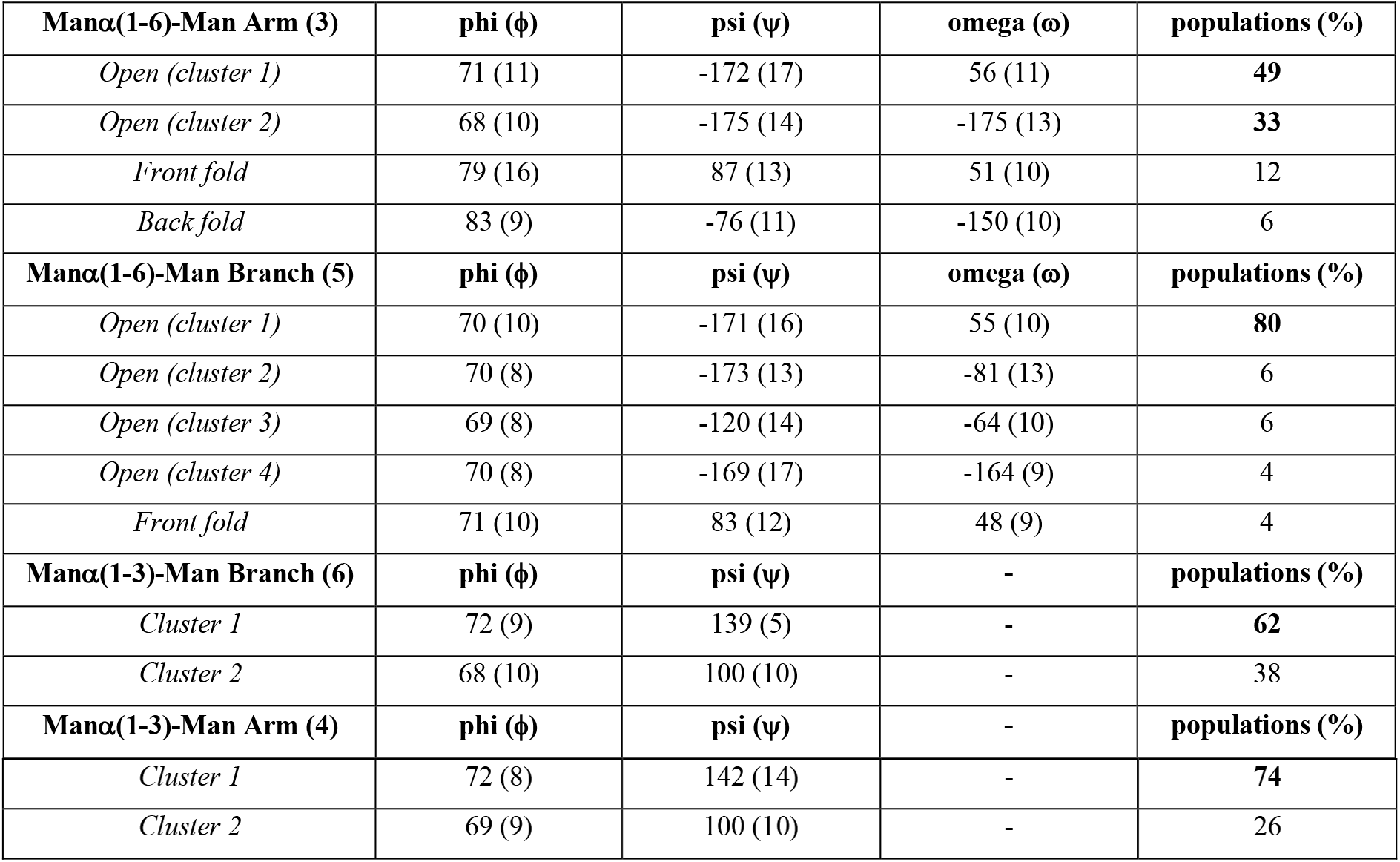
Torsion angles median values of the linkages in the Man5 (1-3/6) arms. Standard deviation values are shown in parenthesis, with relative populations obtained from clustering analysis. Angle values are in degrees (°). The number in parenthesis in the first column indicates the linkages, as shown on the Man5 sketch in **Figure 2**.

**Figure 2.**
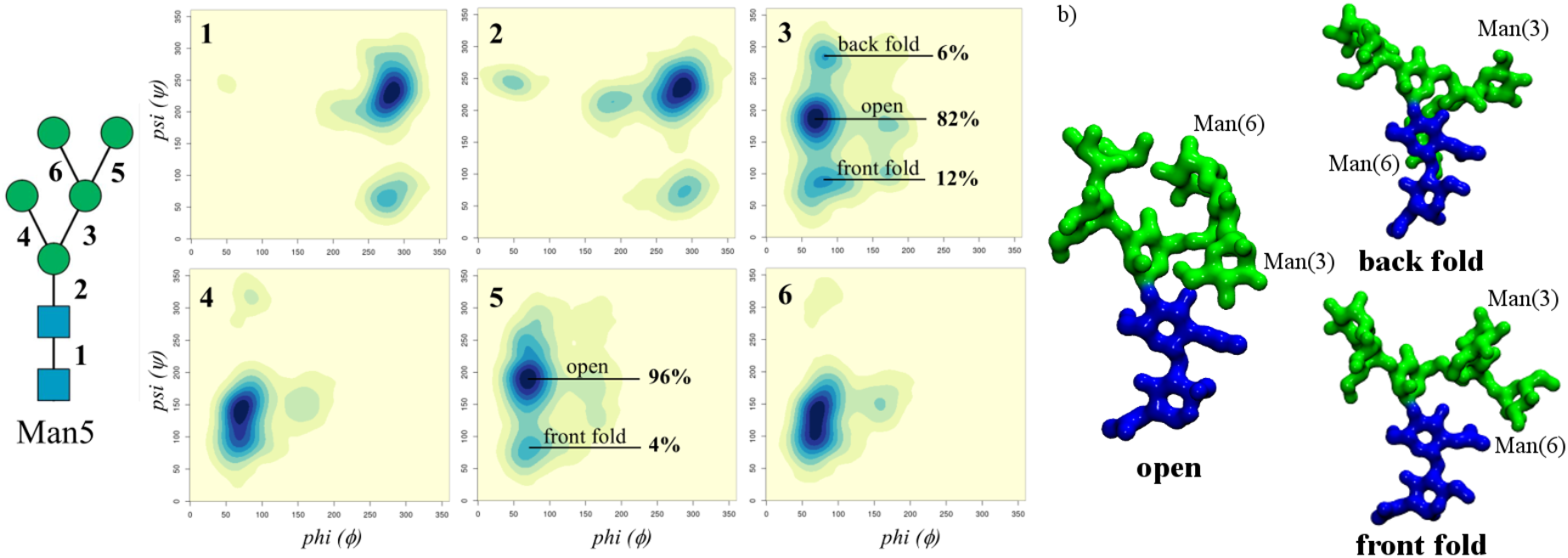
**Panel a)** Man5 conformational analysis in terms of the phi (ϕ) and psi (ψ) torsion angles, with axes ranging from 0 to 360°. Each torsion is numbered as indicated on the left-hand side with the heat maps are labelled on the top-left corner accordingly. **Panel b)** 3D structures of the dominant conformers determined by the flexibility of the (1-6) arm. The Man(3/6) labels indicate the position of the Man on the 3/6 branch on the (1-6) arm. Heat maps made with *RStudio* (www.rstudio.com) and molecular models rendered with *VMD* (www.vmd.org). N-glycans are coloured according to the SNFG convention.

**Figure 3.**
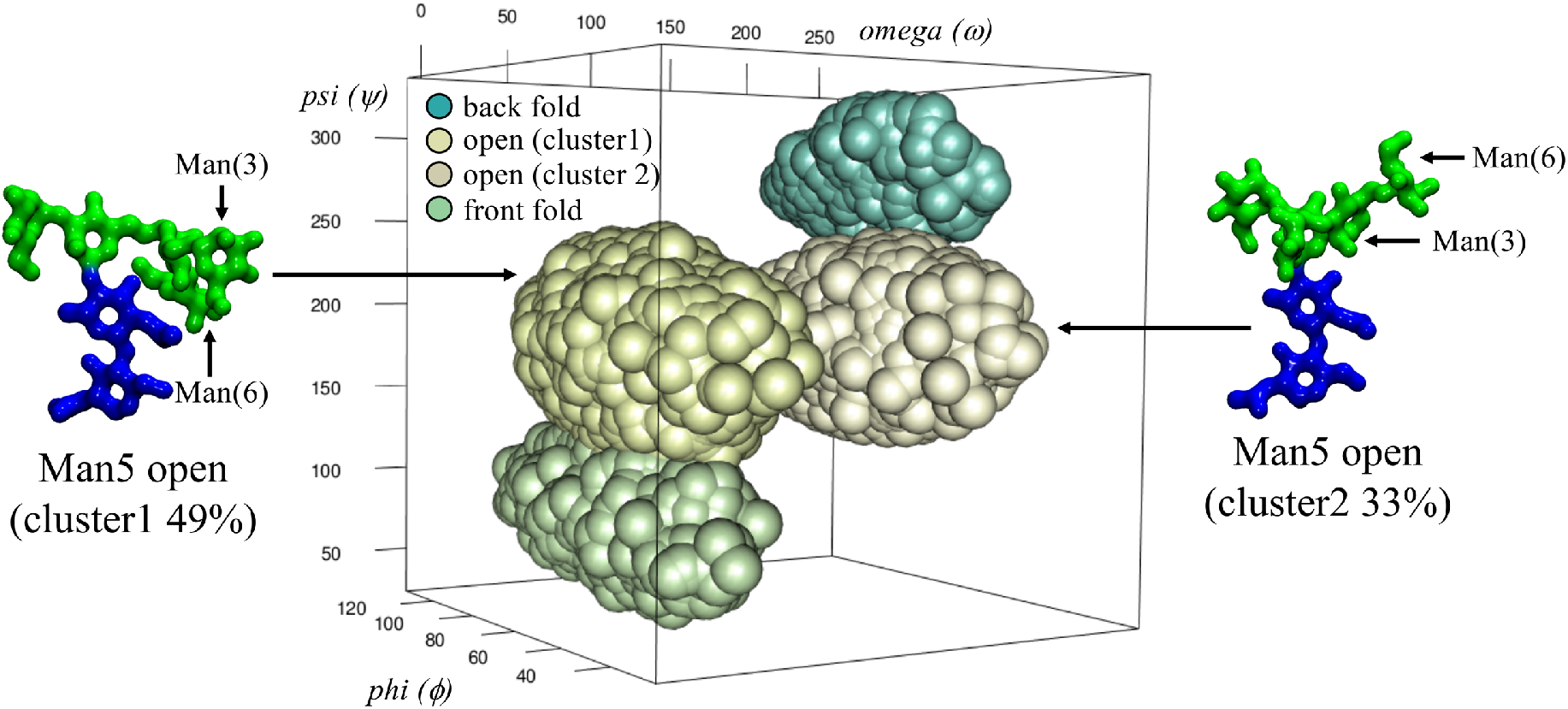
3D representation of the clustering analysis of the Man5 (1-6) arm torsion angles populations. While the back and front fold conformers have one set of values each for the phi, psi and omega torsions, the open conformation adopts two distinct orientations of the biantennary branch, namely open (cluster1) with a representative structure shown on the left-hand side and open (cluster2) with a representative structure shown on the right-hand side. The relative position of the Man(3)- and Man(6)-linked units is also indicated. The *rgl* package in *RStudio* (www.rstudio.com) was used to make the graphics, molecular models rendered with *VMD* (www.vmd.org). N-glycans coloured according to the SNFG convention.

### Man6(I) and Man7(I)

both have a longer (1-3) arm relative to Man5 with one and two Manα (1-2)-Man additional linkages, respectively. Note, different Man6/7 positional isomers exist, where the terminating Man can functionalize either branches on the (1-6) arm. We decided to highlight the Man6/7(I) positional isomers to present the effect of the elongation of the (1-3) arm in combination with a shorter (1-6) arm on the dynamics of the system and their role in enhancing contacts between the arms. A full set of all positional isomers is presented in the Supplementary Material for completeness. As shown in **Figure 4** and **Tables S.2** and **S.6**, both Manα (1-2)-Man linkages occupy two conformers, one at (phi = 74°, psi = 151°) and the other at (phi = 70°, psi = 107°) with a relative populations of 73% and 27% for Man 6, respectively, and one at (phi = 74°, psi = 151°) and the other at (phi = 70°, psi = 106°) with populations of 76% and 24% for Man 7, respectively. As shown by the population analysis in **Tables S.2** and **S.7**, the elongation of the (1-3) arm with Manα (1-2)-Man linkages does not affect the conformational propensity of the (1-6) arm relative to Man5, yet it slightly enhances the flexibility of the (1-3) arm, decreasing the population of the dominant conformer (phi = 72°, psi = 142°) at 74% in Man5, down to 63% in Man7. Notably, the progressive elongation of the (1-3) arm with rigid Manα (1-2)-Man linkages determines an increase of the inter-arms contacts with both (1-6) branches relative to Man5, as discussed in the next subsection. These contacts are stabilized by a complex network of short-lived and interchanging hydrogen bonds that mostly involve the terminal residues of the arms. The open (cluster 1) conformation with α (1-6) torsion values (phi = 71°, psi = -172°, omega =56°) populated at 41% and 46% in Man6 and Man7, respectively, favours the formation of these arm-arm interactions, see **Figure 6 panel b**. Notably, elongation of the (1-3) branch on the (1-6) arm in Man6 (II) and Man 7 (II) determines an increase of these arm-arm interactions that contributes to increasing the population of a previously negligibly populated ‘cluster 3’ conformer, see **Figures S.4** and **S.7** and **Tables S.3** and **S.6**.

**Figure 4.**
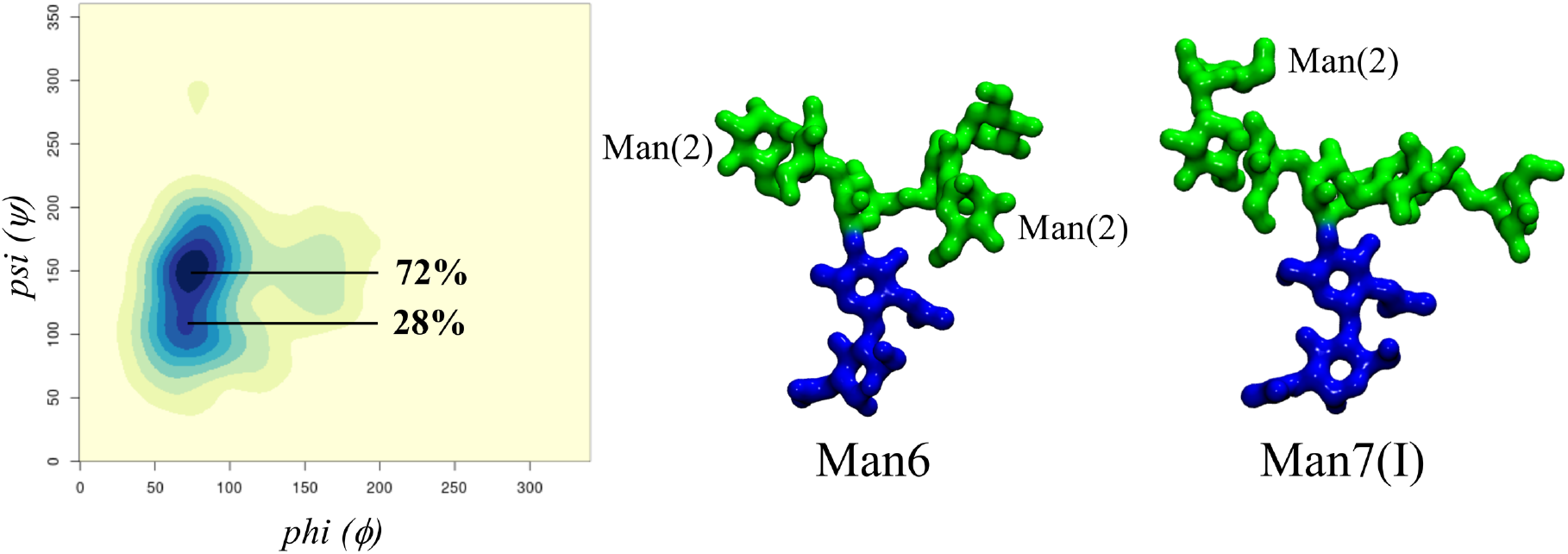
On the left-hand side, heat map representing the conformational analysis with corresponding populations of the first Manα (1-2)-Man linkage on the Man7(I) (1-3) arm, also representative of the corresponding linkage in the Man6 (1-3) arm. On the right-hand side, representative structures of Man6(I) and Man7(I) corresponding to the highest populated Manα (1-2)-Man linkage rotamers in the open (cluster 2) (phi = 71°, psi = -172°, omega =-176°) conformation where the arms do not interact and the conformation of the Manα (1-2)-Man is more clearly visible. Heat maps made with *RStudio* (www.rstudio.com) and molecular models rendered with *VMD* (www.vmd.org). N-glycans coloured according to the SNFG convention.

### Man8(I) and Man9

have a further functionalizations of the (1-6) arm with one Manα (1-2)-Man linkage on the (1-3) branch for the Man8(I) positional isomer and an additional one on (1-6) branch in Man9, see **Figure 1**. As seen for the other oligomannoses, in Man8(I) and Man9 the dominant conformation is with an open (clusters 1, 2 and 3) (1-6) arm, see **Table 2** and **Tables S.9** and **S.13**, with a slightly more pronounced preference for the back *vs* front fold in Man9, due to the interactions of the longer (1-6) branch with the chitobiose, see **Figure 5** and **Table 2**.

**Table 2.** Torsion angles median values of the linkages in the Man9 (1-3/6) arms. Standard deviation values are shown in parenthesis, with relative populations obtained from clustering analysis. Angle values are in degrees (°). The number in parenthesis in the first column indicates the linkages, as shown on the Man9 sketch in **Figure 5**.

| $\text{Man}\alpha(1-6)\text{-Man}$ Arm (3) | $\phi$ ($^\circ$) | $\psi$ ($^\circ$) | $\omega$ ($^\circ$) | populations (%) |
| --- | --- | --- | --- | --- |
| *Open (cluster 1)* | 79 (13) | -173 (16) | 55 (12) | 38 |
| *Open (cluster 2)* | 66 (10) | -179 (13) | -177 (12) | **37** |
| *Open (cluster 3)* | 73 (9) | -157 (21) | -65 (11) | **7** |
| *Cluster 4* | 147 (9) | -172 (8) | 47 (7) | **1** |
| *Front fold* | 78 (11) | 87 (11) | 46 (9) | 3 |
| *Back fold (cluster 1)* | 82 (8) | -74 (10) | -151 (10) | 12 |
| *Back fold (cluster 2)* | 76 (9) | -96 (10) | -75 (9) | 2 |
| **Manα(1-6)-Man Branch (5)** | **phi ($\phi$)** | **psi ($\psi$)** | **omega ($\omega$)** | **populations (%)** |
| *Open (cluster 1)* | 71 (10) | -169 (17) | 55 (10) | **57** |
| *Open (cluster 2)* | 69 (10) | -179 (23) | -166 (11) | **8** |
| *Open (cluster 3)* | 71 (7) | -180 (15) | -73 (8) | **4** |
| *Front fold (Cluster 1)* | 73 (12) | 83 (12) | -164 (18) | 10 |
| *Front fold (Cluster 2)* | 79 (16) | 86 (12) | 51 (10) | 10 |
| *Back fold* | 70 (7) | -118 (12) | -67 (10) | 12 |
| **Manα(1-3)-Man Branch (6)** | **phi ($\phi$)** | **psi ($\psi$)** | - | **populations (%)** |
| *Cluster 1* | 76 (12) | 139 (5) | - | **55** |
| *Cluster 2* | 70 (11) | 100 (10) | - | 26 |
| *Cluster 3* | 148 (11) | 151 (10) | - | 18 |
| **Manα(1-3)-Man Arm (4)** | **phi ($\phi$)** | **psi ($\psi$)** | - | **populations (%)** |
| *Cluster 1* | 72 (9) | 142 (15) | - | **59** |
| *Cluster 2* | 71 (11) | 97 (11) | - | 41 |

**Figure 5.**
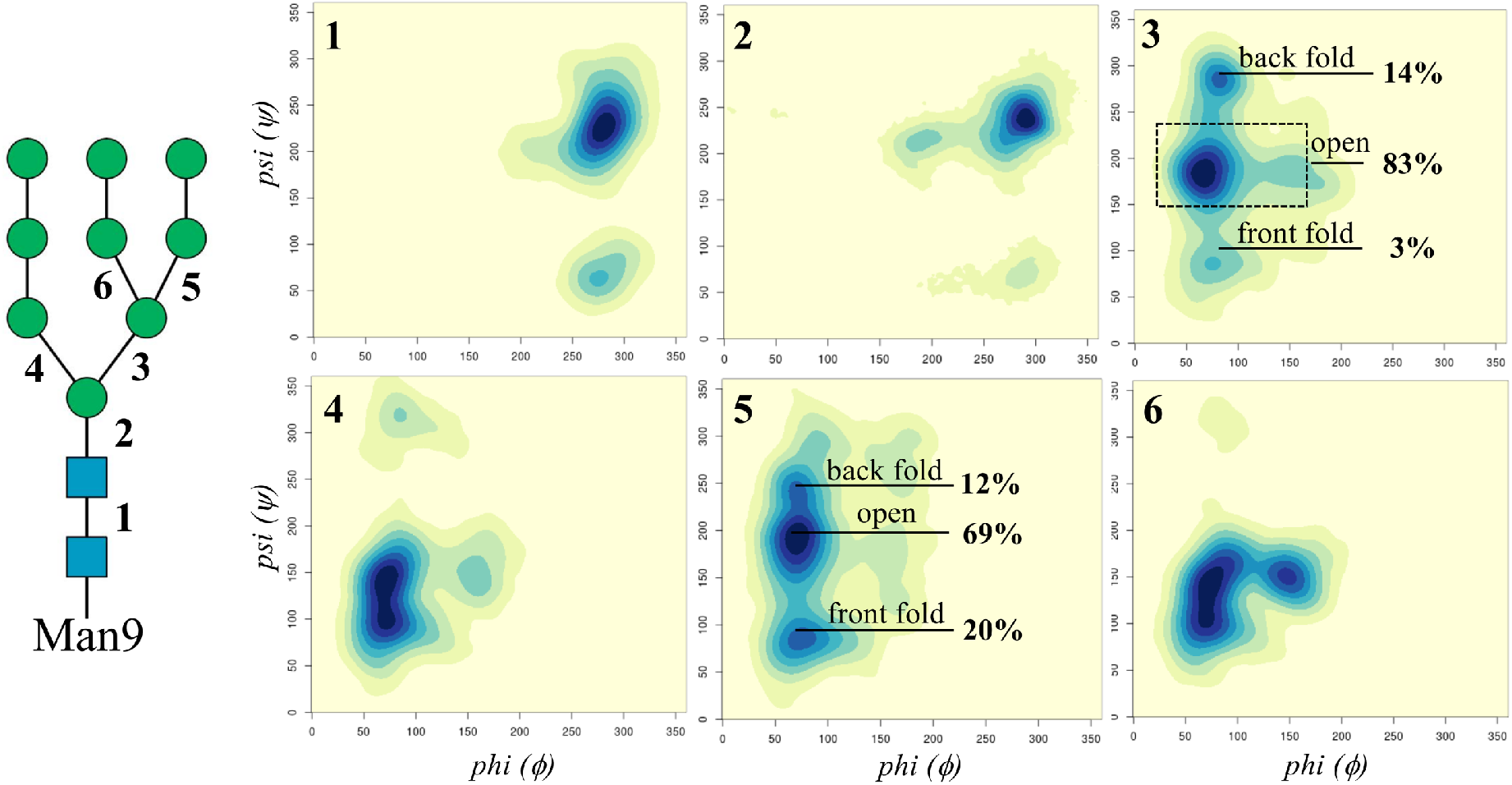
Man9 conformational analysis in terms of the phi (ϕ) and psi (ψ) torsion angles (ranging between 0 and 360°) explored during the 3 μs of cumulative MD sampling. Each torsion was numbered as indicated on the left-hand side sketch and the heat maps have been labelled accordingly on the top left corner. Heat maps were made with *RStudio* (www.rstudio.com).

The structure of all Manα (1-2)-Man linkages is the same as described for Man6(I) and Man7(I), yet the elongation of both branches on the (1-6) arm with relatively rigid Manα (1-2)-Man linkages determines structures with a high number of contacts between the two arms. Indeed, as shown in **Figure 6**, inter-arm contacts only occur within one conformational cluster in Man6(I) and Man7(I), namely open (cluster 1), meanwhile in Man9 interactions between the arms are a feature of virtually all structural populations. These contacts are stabilized by complex networks of rapidly interchanging hydrogen bonds involving mainly the terminal monosaccharides on the arms and branches.

**Figure 6.**
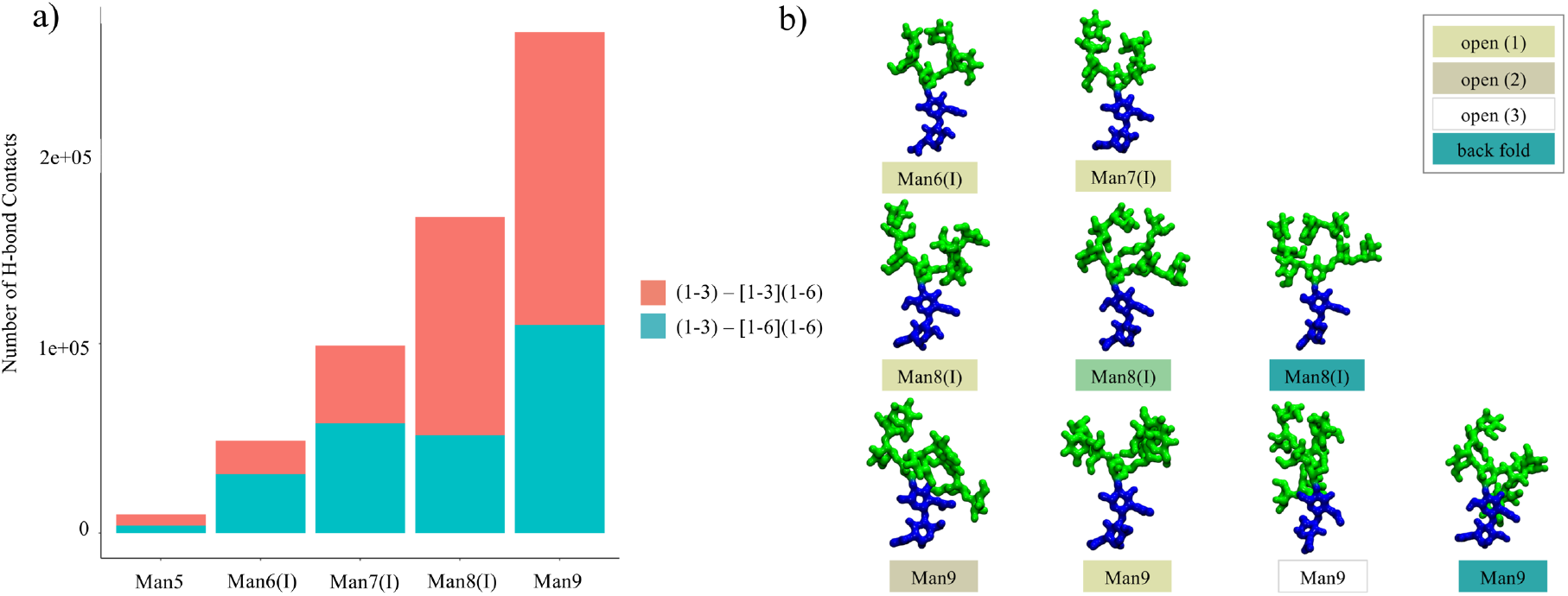
**Panel a)** Number of hydrogen bond contacts (distance threshold 4 Å between donor and acceptor atoms) counted over the 4.5 μs cumulative sampling for each oligomannose indicated on the x-axis. Contacts between the (1-3) arm and the (1-6) branch of the (1-6) arm are shown in cyan. **Panel b)** Representative snapshots from the MD simulations illustrating examples of the inter-arm contacts occurring within each conformational cluster. Different clusters are indicated by the colours in the legend on the top right-hand side, in agreement with the colouring scheme used in Figure 3. Histograms made with *RStudio* (www.rstudio.com) and molecular models rendered with *VMD* (www.vmd.org). N-glycans coloured according to the SNFG convention.

### FcγRIIIa-linked Man5/9

The FcγRIIIa (CD16a) is a cell-bound receptor responsible for modulating antibody-dependent cellular cytotoxicity (ADCC) through its interaction with the IgG1 Fc region^4^. Recent studies have shown that the FcγRIIIa glycosylation contributes to the binding to IgG1s by stabilizing the interaction to a degree that is highly dependent on the type of the N-glycans present^31, 40, 41^. Human FcγRIIIa are glycosylated on two sites, namely N45 and N162, see **Figure 1**. These two sites are very different in terms of their surrounding protein landscape; while N162 is highly exposed to the solvent, N45 is located in the core of one of the two structural domains. To understand the effect of the protein surface landscape on the oligomannoses structure and dynamics, we studied two FcγRIIIa glycoforms, one with Man5 at N45 and N162 and the other with Man9 at N45 and N162.

As shown in **Figure 7**, results obtained from 2 μs of cumulative sampling from two independent runs, show that the conformational dynamics of the Man5 at N45 is significantly restrained compared to the unlinked form. Indeed, a network of hydrogen bonds connects the terminal Man on the (1-3) branch of the (1-6) arm within a protein’s cleft located between the two domains. These interactions result in shifting the Man5 intrinsic conformational equilibrium so that at N45 the Man5 (1-6) arm is mainly allowed in the open (cluster 2) conformation, see **Figure 7** and **Table 3**. The flexibility of the (1-6) branch and of the (1-3) arm, not interacting with the protein, is the same as found for the unlinked Man5, see also **Figure 2**. Man9 has two Manα (1-2)-Man linkages elongating both branches on the (1-6) arm, denying the pose found for Man5, which indeed disappears, see **Tables 3** and **S.13**. Despite a higher flexibility relative to Man5, the N45-linked Man9 is less dynamic relative to the unlinked form due to the protein’s landscape. Indeed, as shown in **Table 3**, only three out of the seven populated conformers are accessible.

**Table 3.** Torsion angles median values of the linkages in the N45-linked Man5 and Man9 (1-6) arm. Standard deviation values are shown in parenthesis, with relative populations obtained from clustering analysis. Angle values are in degrees (°). The number in parenthesis in the first column indicates the linkages, as shown on the Man5 and Man9 sketches in **Figures 2** and **5**.

| N45-Man5 |  |  |  |  |
| --- | --- | --- | --- | --- |
| Manα(1-6)-Man Arm (3) | phi ($\phi$) | psi ($\psi$) | omega ($\omega$) | populations (%) |
| Open (cluster 2) | 69 (10) | -162 (13) | -172 (11) | 100 |
| N45-Man9 |  |  |  |  |
| Open (cluster 1) | 71 (12) | 174 (13) | 59 (15) | 57 |
| Open (cluster 3) | 72 (9) | -174 (15) | -75 (11) | 15 |
| *Back fold (cluster 2)* | 67 (7) | -112 (12) | -69 (12) | 28 |

**Figure 7.**
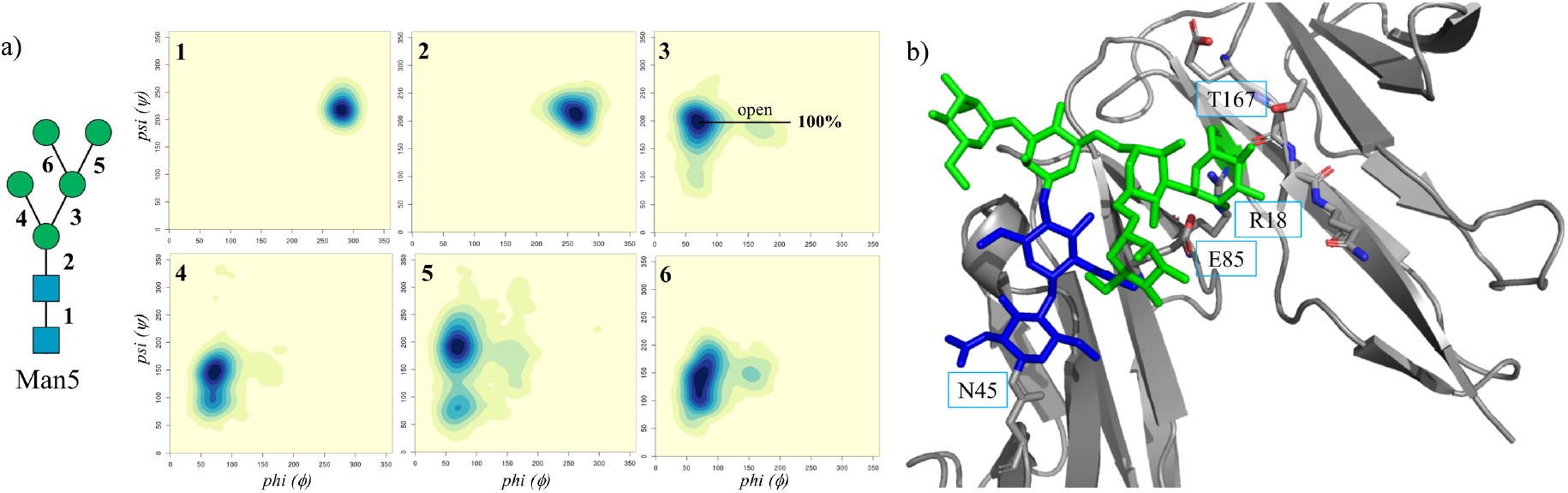
**Panel a)** Conformational analysis in terms of the phi (ϕ) and psi (ψ) torsion angles of the N40-linked Man5 linked explored during the 2 μs of cumulative MD sampling of the Man5 glycosylated FcγRIIIa. Each torsion is numbered as indicated on the left-hand side and the corresponding heat maps are labelled on the top-left corner accordingly. **Panel b)** Dominant conformation of the N45-linked Man5, see **Table 3**, with the terminal man on the (1-3) branch of the (1-6) arm restrained by hydrogen bonds to residues T167, R18 and E85, labelled in the figure. Heat maps made with *RStudio* (www.rstudio.com) and structure rendered with pyMol (www.pymol.org). N-glycan coloured according to the SNFG convention.

As shown in **Figure 1**, the N162 position is much more exposed to the solvent relative to N45. Consequently, the intrinsic dynamics of the N162-Man5 is almost entirely retained, with a shift promoting the open (cluster 2) relative to the open (cluster 1) as the dominant conformer, see **Table 4**. Meanwhile in case of a N162-linked Man9, the dynamics of the longer arms is limited due to the proximity to the protein’s surface, see **Figure 8**, and in particular due to the presence of Lys 128, which because of its position denies a number of conformers due to steric hindrance and also potentially stabilizes the open (cluster 1) conformation through a hydrogen bonding interaction with the α (1-6)-linked Man on the (1-6) arm.

**Table 4.**
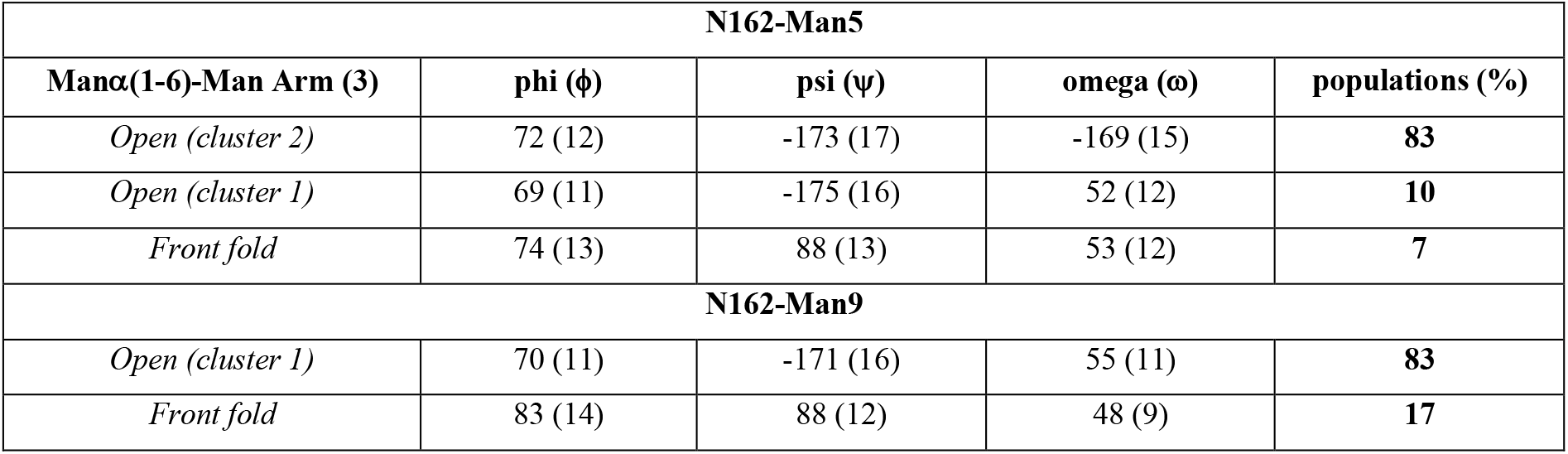
Torsion angles median values of the linkages in the N162-linked Man5 and Man9 (1-6) arm. Standard deviation values are shown in parenthesis, with relative populations obtained from clustering analysis. Angle values are in degrees (°). The number in parenthesis in the first column indicates the linkages, as shown on the Man5 and Man9 sketches in **Figures 2** and **5**.

| N162-Man5 |  |  |  |  |
| --- | --- | --- | --- | --- |
| Man$\alpha(1-6)$-Man Arm (3) | phi ($\phi$) | psi ($\psi$) | omega ($\omega$) | populations (%) |
| *Open (cluster 2)* | 72 (12) | -173 (17) | -169 (15) | **83** |
| *Open (cluster 1)* | 69 (11) | -175 (16) | 52 (12) | **10** |
| *Front fold* | 74 (13) | 88 (13) | 53 (12) | **7** |
| N162-Man9 |  |  |  |  |
| *Open (cluster 1)* | 70 (11) | -171 (16) | 55 (11) | **83** |
| *Front fold* | 83 (14) | 88 (12) | 48 (9) | **17** |

**Figure 8.**
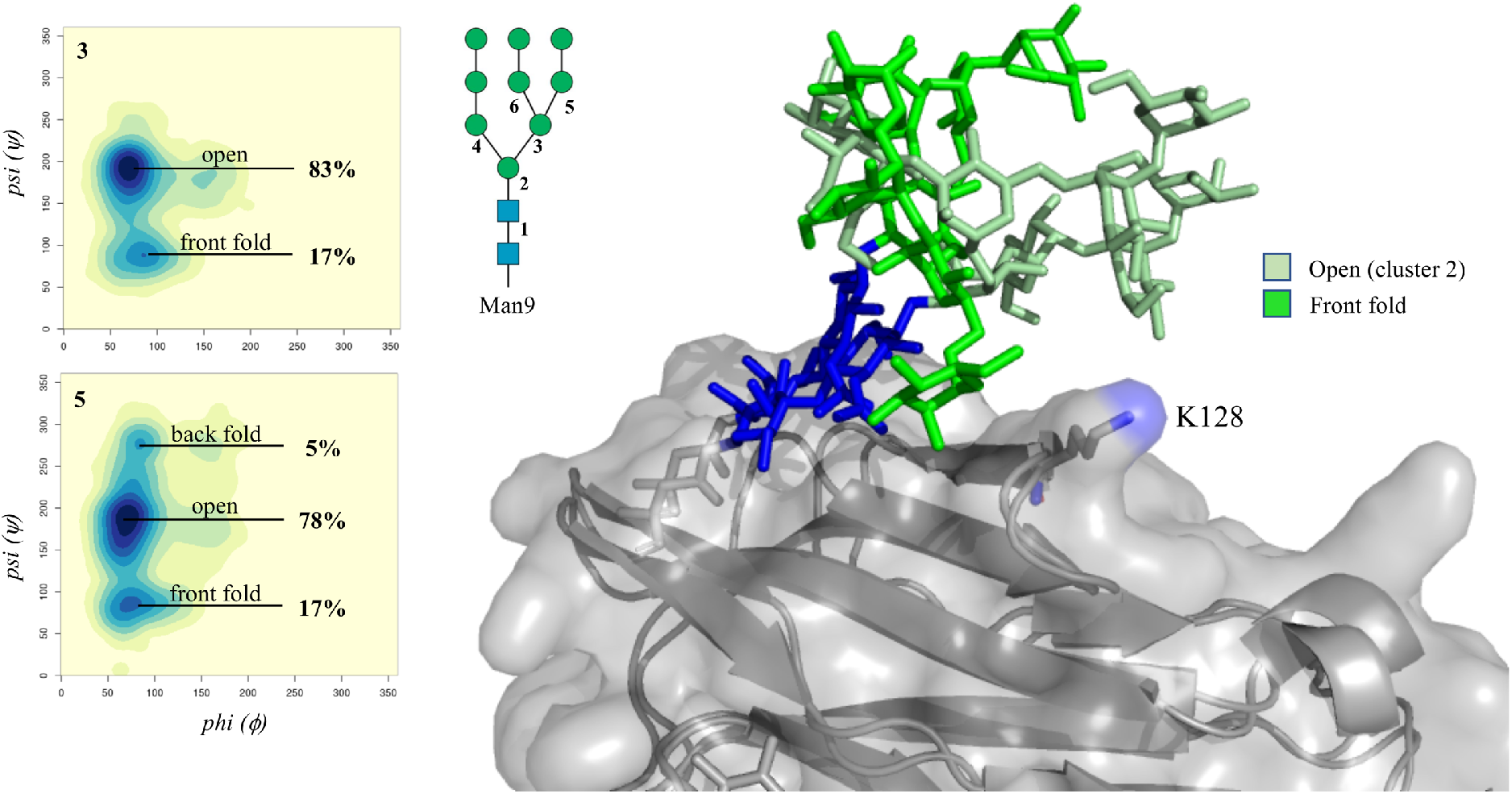
Conformational analysis of the (1-6) arm and (1-6) branch of the N162-linked Man9 in terms of phi (ϕ) and psi (ψ) torsion angles, obtained from the 2 μs of cumulative MD sampling of the Man9 glycosylated FcγRIIIa. Heat maps are labelled on the top-left corner according to the Man9 numbering in the sketch. The two dominant conformations of the N162-linked Man9 are shown on the right-hand side, with the protein represented by the solvent accessible surface and underlying cartoons in grey and the mannose residues with different shades of green as described in the legend. Heat maps made with RStudio (www.rstudio.com) and structure rendered with pyMol (www.pymol.org). N-glycan coloured according to the SNFG convention.

## Discussion

In this work we analysed the 3D structure and dynamics of human oligomannose N-glycans, from Man5 to Man9, when free (unlinked) in solution and also determined how the effect of FcγRIIIa (CD16a) surface landscape modulates their structural equilibria. Despite similarities with complex N-glycans^16, 17^, both in terms of the core chitobiose rigidity and of the relatively low degree of flexibility of the (1-3) arm, oligomannoses have a very unique architecture, which changes with the progressive functionalization of the arms. More specifically, Man5 shows a clear propensity for an ‘open’ structure, where the (1-6) arm is outstretched orientating the two branches on either side of the (1-3) arm, see **Figures 2** and **3**. Small variations of the open structure, determined by two accessible values of the (1-6) arm omega torsion are also populated, see **Figure 3**, and of the additional degrees of freedom of both (1-3/6) branches, which closely reflect the dynamics of the arms, see **Table 1**. In larger oligomannoses the (1-3) arm and (1-3/6) branches on the (1-6) arm are terminated with Manα (1-2)-Man groups, giving raise to different Man6 up to Man8 isoforms and then to Man9. We analysed 10 different positional isomers of Man6 to Man8, see **Figure S.1**, and focused our attention on the isoforms named (I) as representative examples, shown in **Figure 1**, while included all others for completeness as Supplementary Material. Sampling results show that the Manα (1-2)-Man linkages are rigid and do not significantly affect the intrinsic dynamics of the other linkages. The most interesting and unique aspect of the arms elongation in oligomannoses is that the orientation of the additional Manα (1-2)-Man linkages within the underlying architecture of Man5, determines a progressive increase of inter-arm contacts, see **Figure 6**; so, the structure of Man9 is quite compact, or more ‘tree-like’, relative to smaller oligomannoses, where the arms are shorter, but characterized by a more independent dynamics. As a further step in the analysis, a direct comparison of the results we obtained for Man9 and Man8(II) with NMR-validated REMD analysis^42^, show a very good agreement, supporting that the trimming of terminal residues allows for more extended arm structures, which expose embedded glycotopes, see **Figure S.18**.

The results obtained for the unlinked oligomannoses also confirm an earlier observation we made in the context of complex N-glycans^16^, whereby the overall 3D architecture is determined by the local spatial arrangement of independent groups of monosaccharides, we named “glycoblocks”. The oligomannoses dynamics can be also discretized in terms of these structural units^16^, with the addition of a unique Manα (1-2)-Man glycoblock that can be added to the arms, with as we have seen, minimal effect to the dynamics of the underlying units it builds on. This observation can offer a practical advantage to the study of glycan recognition through molecular docking for example, where the receptor binds a specific glycoblock unit and recognition depends only on its accessibility within a specific glycoform.

To understand how the protein affects the presentation of the glycans to potential receptors we have looked at the human FcγRIIIa (CD16a). Human FcγRIIIa has two N-glycosylation sites, namely N45 and N162, where the type of glycosylation affects the receptor’s binding affinity to IgG1s^31, 32, 43^. The surface landscape around these two sites is quite different, with N162 exposed to the solvent, while N45 located in the core of one of the two structural domains, see **Figure 1**. Conformational sampling of a Man5 at N45 shows that the (1-6) arm dynamics is heavily restrained to one of its two open conformations accessible in solution, see **Figure 7**. More specifically, we found that the terminal Man on the (1-3) branch is engaged in a network of hydrogen bonding interactions involving a number of residues near the glycosylation site, namely Arg 18, Glu 85 and Thr 167. The stabilization of this glycoform by the FcγRIIIa surface landscape, renders the (1-3) branch on the (1-6) arm virtually inaccessible for further functionalization. This result agrees with recent work highlighting the unique prevalence of hybrid and oligomannose type N-glycans at N45^32, 43^. The N162 position determines very little steric hindrance to the dynamics of Man5, which retains most of the degrees of freedom characterized for the glycan free in solution. Meanwhile, the dynamics of the larger Man9 is greatly affected by the presence of Lys 128, which forces the glycan to adopt only two of the conformations accessible to the unlinked form, see **Table 4** and **Figure 8**. Ultimately, the comparison between the conformational propensity of the unlinked Man5 and Man9 oligomannoses relative to their FcγRIIIa-linked counterparts suggests that the protein landscape affects the glycans structure by shifting their intrinsic conformational equilibria towards forms that complement it, yet it does not actively morph the glycan into un-natural conformers.

## Conclusions

In this work we have characterized the 3D structure and dynamics of human oligomannose N-glycans unlinked and linked to FcγRIIIa through extensive sampling based on conventional MD simulations. The simulations of the unlinked oligomannose N-glycans show a complex architecture that derives from a progressively intricate network of transient hydrogen bonding interactions involving the terminal residues on the arms, all linked through rigid Manα (1-2)-Man glycoblocks. The protein landscape affects the conformational equilibrium of the N-glycans favouring conformations that complement it, but it does not actively distort the oligomannoses’ structure. Indeed, the two FcγRIIIa glycosylation sites studied in this work present different sets of constraints to different glycoforms and accordingly shift each conformational equilibrium specifically. This determines a diverse degree of accessibility of the arms for further functionalization by glycotransferases and glycohydrolases at N45^32, 43^, which has been found to have an unusually high degree of hybrid N-glycoforms, and ultimately exposure of the arms at N162 for contact with the IgG1 Fc N-glycans, which is implicated in modulating ADCC^19, 31, 44, 45^. Work in this direction is currently ongoing in our lab.

## Supporting Information

The accompanying supporting information includes 17 Tables and 18 Figures providing a complete list of all rotamers conformations with relative populations for all oligomannoses isomers studied in this work. Additional data analysis is shown in Figure S.18.

## Acknowledgements

The Irish Centre for High-End Computing (ICHEC) is gratefully acknowledged for generous allocation of computational resources. The Irish Research Council (IRC) is gratefully acknowledged for funding CAF studies through the Government of Ireland Postgraduate Scholarship Programme.

## Electronic Data Sharing

All structures and trajectories will be made available through a database currently under development in our lab. In the meantime, distribution is done based on requests to the corresponding author.

## References

1. Varki, A., Biological roles of glycans. Glycobiology 2017, 27 (1), 3–49.

2. Strasser, R., Plant protein glycosylation. Glycobiology 2016, 26 (9), 926–939.

3. Christiansen, M., Chik, J., Lee, L., Anugraham, M., Abrahams, J., Packer, N., Cell surface protein glycosylation in cancer. Proteomics 2014, 14 (4-5), 525–546.

4. Cobb, B. A., The history of IgG glycosylation and where we are now. Glycobiology 2020, 30 (4), 202–213.

5. Moremen, K. W., Tiemeyer, M., Nairn, A. V., Vertebrate protein glycosylation: diversity, synthesis and function. Nat Rev Mol Cell Biol 2012, 13 (7), 448–62.

6. Schjoldager, K. T., Narimatsu, Y., Joshi, H. J., Clausen, H., Global view of human protein glycosylation pathways and functions. Nat Rev Mol Cell Biol 2020.

7. Gu, J., Taniguchi, N., Regulation of integrin functions by N-glycans. Glycoconj J 2004, 21 (1-2), 9–15.

8. Strasser, R., Biological significance of complex N-glycans in plants and their impact on plant physiology. Front Plant Sci 2014, 5, 363.

9. Paschinger, K., Wilson, I. B. H., Comparisons of N-glycans across invertebrate phyla. Parasitology 2019, 146 (14), 1733–1742.

10. Deshpande, N., Wilkins, M. R., Packer, N., Nevalainen, H., Protein glycosylation pathways in filamentous fungi. Glycobiology 2008, 18 (8), 626–37.

11. Thompson, A. J., de Vries, R. P., Paulson, J. C., Virus recognition of glycan receptors. Curr Opin Virol 2019, 34, 117–129.

12. Aebi, M., Bernasconi, R., Clerc, S., Molinari, M., N-glycan structures: recognition and processing in the ER. Trends Biochem Sci 2010, 35 (2), 74–82.

13. Rillahan, C. D., Paulson, J. C., Glycan microarrays for decoding the glycome. Annu Rev Biochem 2011, 80, 797–823.

14. Wu, X., Delbianco, M., Anggara, K., Michnowicz, T., Pardo-Vargas, A., Bharate, P., Sen, S., Pristl, M., Rauschenbach, S., Schlickum, U. et al.., Imaging single glycans. Nature 2020, 582 (7812), 375–378.

15. Anggara, K., Zhu, Y., Delbianco, M., Rauschenbach, S., Abb, S., Seeberger, P. H., Kern, K., Exploring the Molecular Conformation Space by Soft Molecule-Surface Collision. J Am Chem Soc 2020.

16. Fogarty, C. A., Harbison, A. M., Dugdale, A. R., Fadda, E., How and why plants and human N-glycans are different: Insight from molecular dynamics into the “glycoblocks” architecture of complex carbohydrates. Beilstein J Org Chem 2020, 16, 2046–2056.

17. Harbison, A. M., Brosnan, L. P., Fenlon, K., Fadda, E., Sequence-to-structure dependence of isolated IgG Fc complex biantennary N-glycans: a molecular dynamics study. Glycobiology 2019, 29 (1), 94–103.

18. Neelamegham, S., Aoki-Kinoshita, K., Bolton, E., Frank, M., Lisacek, F., Lütteke, T., O’Boyle, N., Packer, N. H., Stanley, P., Toukach, P. et al.., Group, S. D., Updates to the Symbol Nomenclature for Glycans guidelines. Glycobiology 2019, 29 (9), 620–624.

19. Ferrara, C., Grau, S., Jäger, C., Sondermann, P., Brünker, P., Waldhauer, I., Hennig, M., Ruf, A., Rufer, A. C., Stihle, M., Umaña, P., Benz, J., Unique carbohydrate-carbohydrate interactions are required for high affinity binding between FcgammaRIII and antibodies lacking core fucose. Proc Natl Acad Sci U S A 2011, 108 (31), 12669–74.

20. Watanabe, Y., Bowden, T. A., Wilson, I. A., Crispin, M., Exploitation of glycosylation in enveloped virus pathobiology. Biochim Biophys Acta Gen Subj 2019, 1863 (10), 1480–1497.

21. Watanabe, Y., Allen, J. D., Wrapp, D., McLellan, J. S., Crispin, M., Site-specific glycan analysis of the SARS-CoV-2 spike. Science 2020.

22. Bonomelli, C., Doores, K. J., Dunlop, D. C., Thaney, V., Dwek, R. A., Burton, D. R., Crispin, M., Scanlan, C. N., The glycan shield of HIV is predominantly oligomannose independently of production system or viral clade. PLoS One 2011, 6 (8), e23521.

23. Doores, K. J., Bonomelli, C., Harvey, D. J., Vasiljevic, S., Dwek, R. A., Burton, D. R., Crispin, M., Scanlan, C. N., Envelope glycans of immunodeficiency virions are almost entirely oligomannose antigens. Proc Natl Acad Sci U S A 2010, 107 (31), 13800–5.

24. Stewart-Jones, G., Soto, C., Lemmin, T., Chuang, G., Druz, A., Kong, R., Thomas, P., Wagh, K., Zhou, T., Behrens, A. et al.., l Trimeric HIV-1-Env Structures Define Glycan Shields from Clades A, B, and G. Cell 2016, 165 (4), 813–826.

25. Struwe, W. B., Chertova, E., Allen, J. D., Seabright, G. E., Watanabe, Y., Harvey, D. J., Medina-Ramirez, M., Roser, J. D., Smith, R., Westcott, D. et al.., Site-Specific Glycosylation of Virion-Derived HIV-1 Env Is Mimicked by a Soluble Trimeric Immunogen. Cell Rep 2018, 24 (8), 1958-1966.e5.

26. de Leoz, M. L., Young, L. J., An, H. J., Kronewitter, S. R., Kim, J., Miyamoto, S., Borowsky, A. D., Chew, H. K., Lebrilla, C. B., High-mannose glycans are elevated during breast cancer progression. Mol Cell Proteomics 2011, 10 (1), M110.002717.

27. Li, Q., Li, G., Zhou, Y., Zhang, X., Sun, M., Jiang, H., Yu, G., Comprehensive N-Glycome Profiling of Cells and Tissues for Breast Cancer Diagnosis. J Proteome Res 2019, 18 (6), 2559–2570.

28. Liu, X., Nie, H., Zhang, Y., Yao, Y., Maitikabili, A., Qu, Y., Shi, S., Chen, C., Li, Y., Cell surface-specific N-glycan profiling in breast cancer. PLoS One 2013, 8 (8), e72704.

29. Thaysen-Andersen, M., Packer, N. H., Site-specific glycoproteomics confirms that protein structure dictates formation of N-glycan type, core fucosylation and branching. Glycobiology 2012, 22 (11), 1440–52.

30. Zacchi, L. F., Schulz, B. L., N-glycoprotein macroheterogeneity: biological implications and proteomic characterization. Glycoconj J 2016, 33 (3), 359–76.

31. Subedi, G. P., Barb, A. W., CD16a with oligomannose-type. J Biol Chem 2018, 293 (43), 16842–16850.

32. Roberts, J. T., Patel, K. R., Barb, A. W., Site-specific N-glycan Analysis of Antibody-binding Fc γ Receptors from Primary Human Monocytes. Mol Cell Proteomics 2020, 19 (2), 362–374.

33. Case, D., Ben-Shalom, I., Brozell, S., Cerutti, D., Cheatham III, T., Cruzeiro, V., Darden, T., Duke, R., Ghoreishi, D., Gilson, M. et al.., AMBER 2018, University of California, San Francisco, 2018.

34. Kirschner, K. N., Yongye, A. B., Tschampel, S. M., González-Outeiriño, J., Daniels, C. R., Foley, B. L., Woods, R. J., GLYCAM06: a generalizable biomolecular force field. Carbohydrates. J Comput Chem 2008, 29 (4), 622–55.

35. Jorgensen, W., Chandrasekhar, J., Madura, J., Impey, R., Klein, M., Comparison of simple potential functions for simulations of liquid water. Journal of Chemical Physics 1983, 79 (2), 926–935.

36. Fadda, E., Woods, R. J., On the Role of Water Models in Quantifying the Binding Free Energy of Highly Conserved Water Molecules in Proteins: The Case of Concanavalin A. J Chem Theory Comput 2011, 7 (10), 3391–8.

37. Sauter, J., Grafmüller, A., Solution Properties of Hemicellulose Polysaccharides with Four Common Carbohydrate Force Fields. J Chem Theory Comput 2015, 11 (4), 1765–74.

38. Joung, I. S., Cheatham, T. E., Determination of alkali and halide monovalent ion parameters for use in explicitly solvated biomolecular simulations. J Phys Chem B 2008, 112 (30), 9020–41.

39. Humphrey, W., Dalke, A., Schulten, K., VMD: visual molecular dynamics. J Mol Graph 1996, 14 (1), 33-8, 27-8.

40. Hayes, J. M., Frostell, A., Karlsson, R., Müller, S., Martín, S. M., Pauers, M., Reuss, F., Cosgrave, E. F., Anneren, C., Davey, G. P., Rudd, P. M., Identification of Fc Gamma Receptor Glycoforms That Produce Differential Binding Kinetics for Rituximab. Mol Cell Proteomics 2017, 16 (10), 1770–1788.

41. Subedi, G. P., Barb, A. W., The Structural Role of Antibody N-Glycosylation in Receptor Interactions. Structure 2015, 23 (9), 1573–1583.

42. Yamaguchi, T., Sakae, Y., Zhang, Y., Yamamoto, S., Okamoto, Y., Kato, K., Exploration of conformational spaces of high-mannose-type oligosaccharides by an NMR-validated simulation. Angew Chem Int Ed Engl 2014, 53 (41), 10941–4.

43. Patel, K. R., Roberts, J. T., Barb, A. W., Allotype-specific processing of the CD16a N45-glycan from primary human natural killer cells and monocytes. Glycobiology 2020, 30 (7), 427–432.

44. Falconer, D. J., Subedi, G. P., Marcella, A. M., Barb, A. W., Antibody Fucosylation Lowers the FcγRIIIa/CD16a Affinity by Limiting the Conformations Sampled by the N162-Glycan. ACS Chem Biol 2018, 13 (8), 2179–2189.

45. Day, C. J., Tran, E. N., Semchenko, E. A., Tram, G., Hartley-Tassell, L. E., Ng, P. S., King, R. M., Ulanovsky, R., McAtamney, S., Apicella, M. A. et al.., Glycan:glycan interactions: High affinity biomolecular interactions that can mediate binding of pathogenic bacteria to host cells. Proc Natl Acad Sci U S A 2015, 112 (52), E7266–75.

