## Supplementary Information for "The oligomannose N-glycans 3D architecture and its response to the FcγRIIIa structural landscape"

#### Contents

**1 Oligomannose Isomers** **2**

**2 Man 5** **3**

**3 Man 6 I** **5**

**4 Man 6 II** **7**

**5 Man 6 III** **9**

**6 Man 7 I** **11**

**7 Man 7 II** **13**

**8 Man 7 III** **15**

**9 Man 7 IV** **17**

**10 Man 8 I** **19**

**11 Man 8 II** **21**

**12 Man 8 III** **23**

**13 Man 9** **25**

**14 Fc$\gamma$RC: Man5 N45** **27**

**15 Fc$\gamma$RC: Man5 N162** **29**

**16 Fc$\gamma$RC: Man9 N45** **31**

**17 Fc$\gamma$RC: Man9 N162** **33**

**18 DBSCAN Parameters** **35**

**19 Man 9 / 8(II) Distance Measurements** **36**

### 1 Oligomannose Isomers

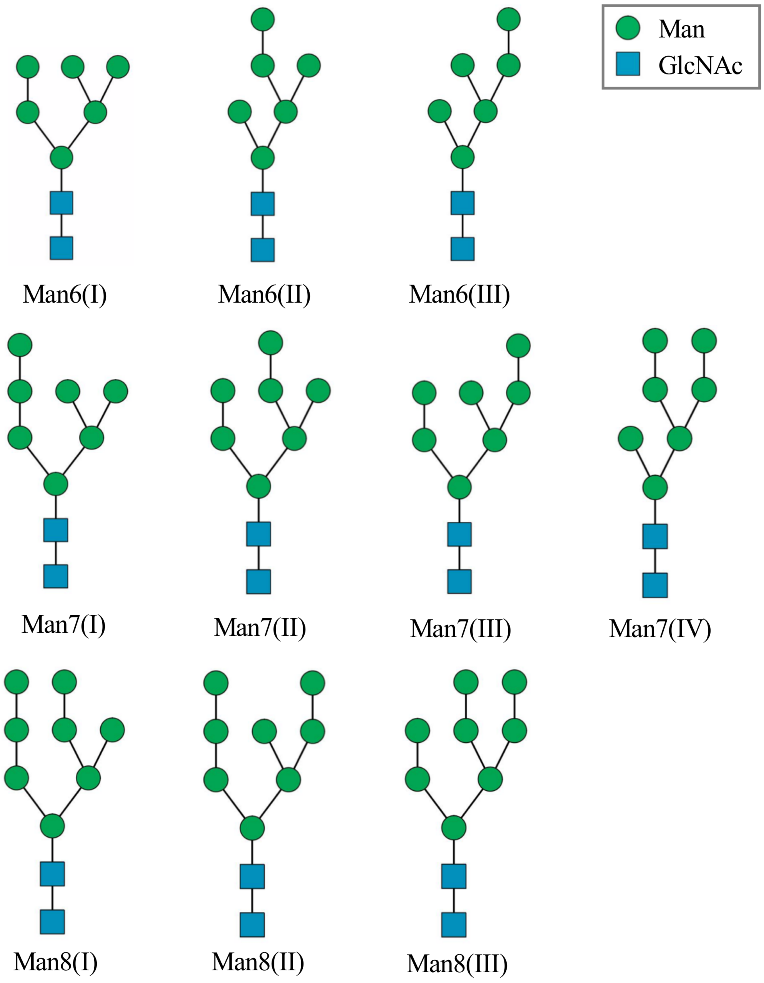


Figure 1: SNFG representation of all the Man6-8 oligomannose positional isomers studied in this work in addition to Man5/9

### 2 Man 5

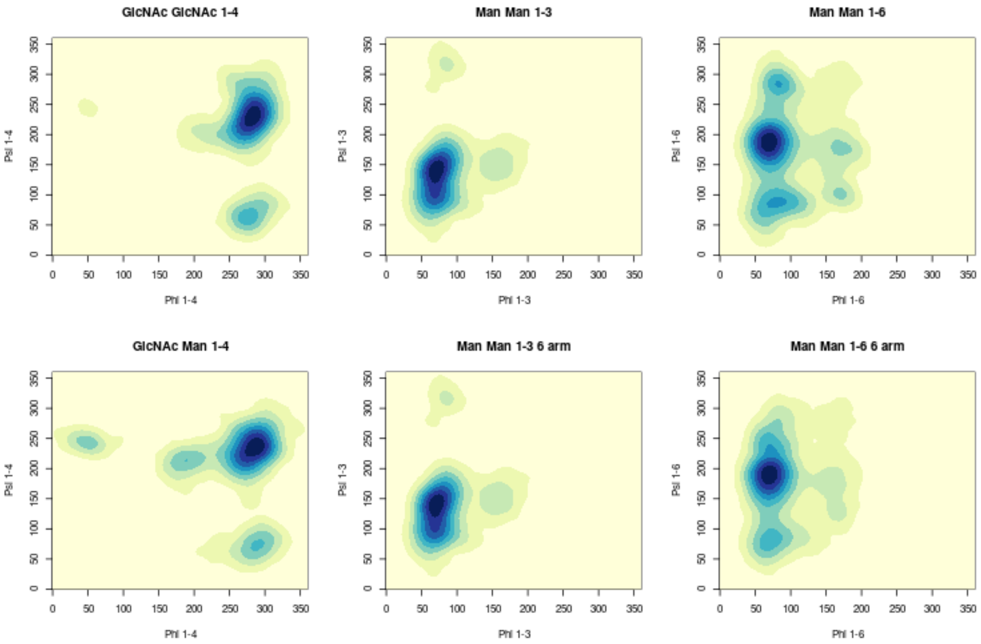


Figure 2: 2D Kernel density estimates for the $\phi$ / $\psi$ angle distributions for Man 5.

Table 1: The $\phi$ / $\psi$ / $\omega$ angle distributions for Man 5.

| **GlcNAc $\beta$(1-4) GlcNAc** | $\phi$ | $\psi$ | $\omega$ | **Pop (%)** |
|---|---|---|---|---|
| Cluster 1 | -78.1 (11.2) | -130.2 (18.3) | - | 94.6 |
| Cluster 2 | -81.9 (11.4) | 65.5 (11.2) | - | 5.4 |
| **Man $\beta$(1-4) GlcNAc** | $\phi$ | $\psi$ | $\omega$ | **Pop (%)** |
| Cluster 1 | -76.0 (12.9) | -125.9 (15.6) | - | 93.0 |
| Cluster 2 | -70.5 (10.7) | 72.9 (10.9) | - | 3.5 |
| Cluster 3 | -170.7 (12.3) | -146.8 (7.83) | - | 2.5 |
| Cluster 4 | 48.6 (12.9) | -116.3 (5.5) | - | 1.0 |
| **Man $\alpha$(1-3) Man (1-3)** | $\phi$ | $\psi$ | $\omega$ | **Pop (%)** |
| Cluster 1 | 71.7 (8.1) | 141.6 (14.5) | - | 73.9 |
| Cluster 2 | 68.8 (9.4) | 99.7 (10.5) | - | 26.1 |
| **Man $\alpha$(1-3) Man (1-6)** | $\phi$ | $\psi$ | $\omega$ | **Pop (%)** |
| Cluster 1 | 72.1 (9.3) | 138.6 (15.1) | - | 62.5 |
| Cluster 2 | 67.6 (9.9) | 99.9 (10.4) | - | 37.5 |
| **Man $\alpha$(1-6) Man** | $\phi$ | $\psi$ | $\omega$ | **Pop (%)** |
| Cluster 1 | 71.1 (10.7) | -172.5 (17.0) | 56.1 (10.8) | 48.6 |
| Cluster 2 | 67.8 (10.0) | -175.4 (14.0) | -175.4 (12.7) | 33.3 |
| Cluster 3 | 79.2 (16.4) | 86.7 (13.4) | 50.9 (10.0) | 12.4 |
| Cluster 4 | 82.6 (8.6) | -76.5 (10.8) | -150.2 (10.5) | 5.7 |
| **Man $\alpha$(1-6) Man (1-6)** | $\phi$ | $\psi$ | $\omega$ | **Pop (%)** |
| Cluster 1 | 70.3 (10.3) | -171.2 (15.9) | 54.8 (10.5) | 80.4 |
| Cluster 2 | 69.6 (8.5) | -173.3 (13.4) | -80.7 (12.6) | 5.7 |
| Cluster 3 | 69.6 (7.6) | -120.2 (13.6) | -64.3 (10.5) | 5.6 |
| Cluster 4 | 69.7 (7.6) | -168.7 (17.5) | -164.4(9.4) | 4.4 |
| Cluster 5 | 70.9 (10.5) | 82.9 (11.6) | 47.9 (8.8) | 3.9 |

### 3 Man 6 I

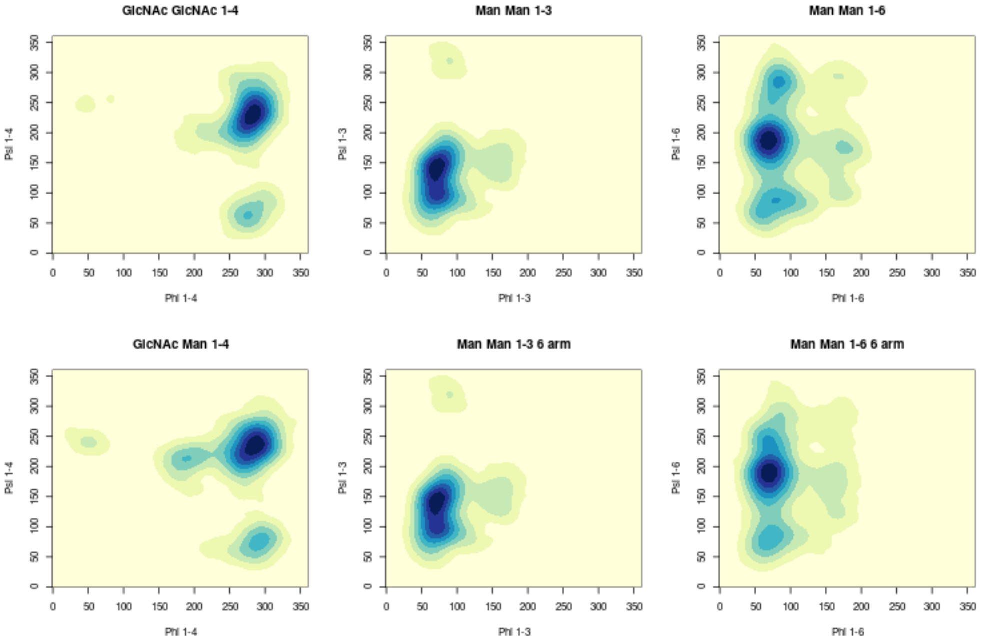


Figure 3: 2D Kernel density estimates for the $\phi$ / $\psi$ angle distributions for Man 6 I.

Table 2: The $\phi$ / $\psi$ / $\omega$ angle distributions for Man 6 I.

| **GlcNAc $\beta$(1-4) GlcNAc** | $\phi$ | $\psi$ | $\omega$ | **Pop (%)** |
|---|---|---|---|---|
| Cluster 1 | -78.0 (10.9) | -130.3 (17.4) | - | 96.7 |
| Cluster 2 | -82.5 (10.7) | 64.7 (11.1) | - | 3.3 |
| **Man $\beta$(1-4) GlcNAc** | $\phi$ | $\psi$ | $\omega$ | **Pop (%)** |
| Cluster 1 | -76.2 (12.9) | -126.4 (15.4) | - | 92.4 |
| Cluster 2 | -68.5 (10.5) | 74.4 (10.94) | - | 4.8 |
| Cluster 3 | -169.1 (12.1) | -146.2 (8.3) | - | 2.8 |
| **Man $\alpha$(1-3) Man (1-3)** | $\phi$ | $\psi$ | $\omega$ | **Pop (%)** |
| Cluster 1 | 72.1 (9.1) | 142.1 (14.4) | - | 67.8 |
| Cluster 2 | 71.1 (9.9) | 98.5 (10.4) | - | 32.2 |
| **Man $\alpha$(1-3) Man (1-6)** | $\phi$ | $\psi$ | $\omega$ | **Pop (%)** |
| Cluster 1 | 72.3 (9.2) | 138.6 (15.1) | - | 62.2 |
| Cluster 2 | 67.7 (9.7) | 99.7 (10.3) | - | 37.8 |
| **Man $\alpha$(1-6) Man** | $\phi$ | $\psi$ | $\omega$ | **Pop (%)** |
| Cluster 1 | 71.0 (10.7) | -172.2 (15.8) | 55.9 (10.8) | 41.1 |
| Cluster 2 | 67.7 (9.9) | -175.1 (14.1) | -175.8 (12.3) | 37.5 |
| Cluster 3 | 76.6 (16.5) | 85.1 (14.6) | 50.3 (9.9) | 10.9 |
| Cluster 4 | 83.3 (8.5) | -75.3 (11.7) | -149.0 (10.8) | 7.1 |
| Cluster 5 | 70.7 (8.7) | -177.6 (13.6) | -71.9 (10.1) | 3.4 |
| **Man $\alpha$(1-6) Man (1-6)** | $\phi$ | $\psi$ | $\omega$ | **Pop (%)** |
| Cluster 1 | 70.2 (10.4) | -171.3 (15.7) | 54.7 (10.3) | 76.0 |
| Cluster 2 | 69.2 (8.7) | -173.9 (13.6) | -82.4 (13.3) | 7.8 |
| Cluster 3 | 68.7 (7.8) | -119.6 (14.1) | -65.1 (10.8) | 8.5 |
| Cluster 4 | 70.2 (7.8) | -175.5 (17.5) | -163.6 (8.9) | 4.4 |
| Cluster 5 | 71.3 (9.3) | 82.0 (10.1) | 47.5 (8.0) | 3.2 |
| **Man $\alpha$(1-2) Man (1-3)** | $\phi$ | $\psi$ | $\omega$ | **Pop (%)** |
| Cluster 1 | 74.3 (8.7) | 150.7 (15.1) | - | 72.7 |
| Cluster 2 | 70.0 (9.4) | 107.2 (11.6) | - | 27.3 |

### 4 Man 6 II

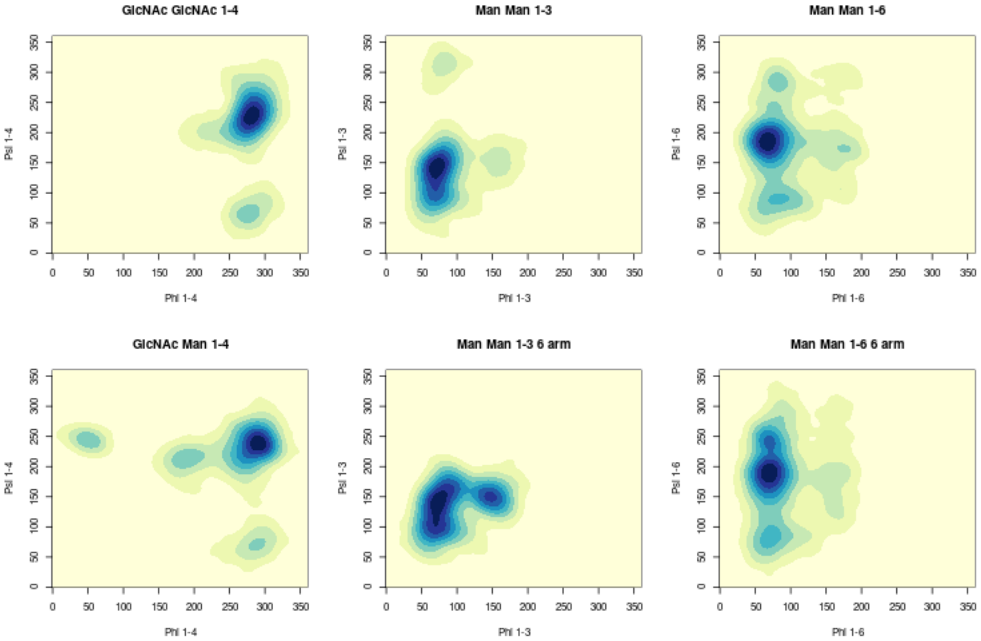


Figure 4: 2D Kernel density estimates for the $\phi$ / $\psi$ angle distributions for Man 6 II.

Table 3: The $\phi$ / $\psi$ / $\omega$ angle distributions for Man 6 II.

| **GlcNAc $\beta$(1-4) GlcNAc** | $\phi$ | $\psi$ | $\omega$ | **Pop (%)** |
|---|---|---|---|---|
| Cluster 1 | -78.9 (10.3) | -131.7 (16.7) | - | 98.0 |
| Cluster 2 | -83.0 (9.4) | 65.8 (9.0) | - | 2.0 |
| **Man $\beta$(1-4) GlcNAc** | $\phi$ | $\psi$ | $\omega$ | **Pop (%)** |
| Cluster 1 | -71.5 (11.8) | -122.8 (13.8) | - | 94.2 |
| Cluster 2 | -70.3 (7.8) | 70.7 (6.5) | - | 1.0 |
| Cluster 3 | -170.2 (12.0) | -146.2 (8.0) | - | 2.8 |
| Cluster 4 | 50.7 (8.2) | -116.0 (5.7) | - | 2.0 |
| **Man $\alpha$(1-3) Man (1-3)** | $\phi$ | $\psi$ | $\omega$ | **Pop (%)** |
| Cluster 1 | 72.3 (9.2) | 138.6 (15.1) | - | 62.2 |
| Cluster 2 | 67.7 (9.7) | 99.7 (10.3) | - | 37.8 |
| **Man $\alpha$(1-3) Man (1-6)** | $\phi$ | $\psi$ | $\omega$ | **Pop (%)** |
| Cluster 1 | 76.1 (11.3) | 144.6 (16.6) | - | 57.0 |
| Cluster 2 | 70.3 (10.5) | 100.3 (9.5) | - | 24.8 |
| Cluster 3 | 147.4 (10.5) | 150.2 (9.6) | - | 18.3 |
| **Man $\alpha$(1-6) Man** | $\phi$ | $\psi$ | $\omega$ | **Pop (%)** |
| Cluster 1 | 71.0 (10.7) | -172.2 (15.8) | 56.2 (11.4) | 58.2 |
| Cluster 2 | 65.4 (16.3) | -178.9 (11.8) | -177.4 (11.2) | 33.9 |
| Cluster 3 | 82.1 (16.4) | 89.2 (13.7) | 51.8 (9.9) | 6.3 |
| Cluster 4 | 81.7 (7.4) | -75.6 (9.1) | -149.2 (9.2) | 1.6 |
| **Man $\alpha$(1-6) Man (1-6)** | $\phi$ | $\psi$ | $\omega$ | **Pop (%)** |
| Cluster 1 | 70.3 (10.5) | -171.4 (16.2) | 54.7 (10.6) | 70.28 |
| Cluster 2 | 71.3(7.1) | 179.1 (10.22) | -72.0 (8.3) | 3.9 |
| Cluster 3 | 69.5 (7.2) | -119.1 (14.0) | -67.1 (9.5) | 14.5 |
| Cluster 4 | 70.7 (9.7) | -173.3 (23.8) | -164.6 (13.2) | 6.9 |
| Cluster 5 | 71.5 (12.0) | 84.2 (13.6) | 49.0 (10.2) | 4.3 |
| **Man $\alpha$(1-2) Man (1-3)** | $\phi$ | $\psi$ | $\omega$ | **Pop (%)** |
| Cluster 1 | 72.3 (8.0) | 149.3 (13.4) | - | 74.8 |
| Cluster 2 | 69.6 (7.9) | 112.5 (10.1) | - | 25.2 |

### 5 Man 6 III

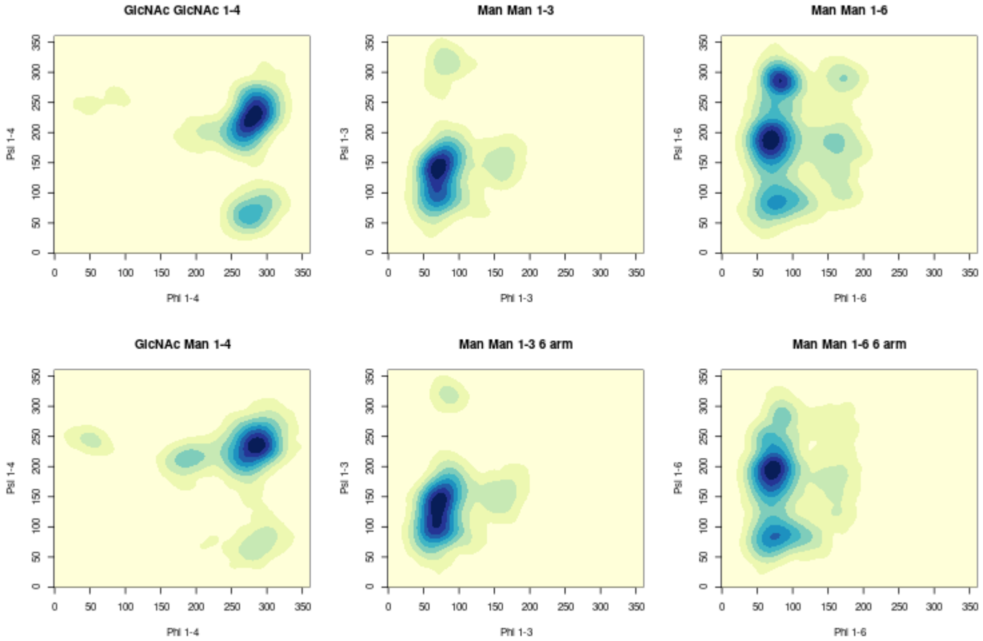


Figure 5: 2D Kernel density estimates for the $\phi$ / $\psi$ angle distributions for Man 6 III.

Table 4: The $\phi$ / $\psi$ / $\omega$ angle distributions for Man 6 III.

| **GlcNAc $\beta$(1-4) GlcNAc** | $\phi$ | $\psi$ | $\omega$ | **Pop (%)** |
|---|---|---|---|---|
| Cluster 1 | -78.8 (11.4) | -133.8 (17.5) | - | 93.5 |
| Cluster 2 | -82.0 (11.8) | 66.0 (11.7) | - | 6.5 |
| **Man $\beta$(1-4) GlcNAc** | $\phi$ | $\psi$ | $\omega$ | **Pop (%)** |
| Cluster 1 | - 76.3 (12.6) | -125.5 (14.4) | - | 97.9 |
| Cluster 2 | -170.4 (10.9) | -146.2 (7.5) | - | 2.1 |
| **Man $\alpha$(1-3) Man (1-3)** | $\phi$ | $\psi$ | $\omega$ | **Pop (%)** |
| Cluster 1 | 76.1 (11.3) | 144.6 (16.6) | - | 57.0 |
| Cluster 2 | 70.3 (10.5) | 100.3 (9.5) | - | 24.8 |
| **Man $\alpha$(1-3) Man (1-6)** | $\phi$ | $\psi$ | $\omega$ | **Pop (%)** |
| Cluster 1 | 72.3 (9.2) | 138.6 (15.1) | - | 62.2 |
| Cluster 2 | 67.7 (9.7) | 99.7 (10.3) | - | 37.8 |
| **Man $\alpha$(1-6) Man** | $\phi$ | $\psi$ | $\omega$ | **Pop (%)** |
| Cluster 1 | 71.2 (10.7) | -172.9 (17.2) | 56.0 (10.7) | 36.1 |
| Cluster 2 | 67.5 (10.7) | -174.3 (15.2) | -175.9 (12.4) | 28.6 |
| Cluster 3 | 79.1 (14.8) | 86.4 (12.9) | 50.1 (10.7) | 9.7 |
| Cluster 4 | 82.3 (7.7) | -74.5 (9.8) | -151.5 (9.7) | 24.2 |
| Cluster 5 | 71.6 (6.9) | -173.2 (11.5) | -69.2 (8.3) | 1.4 |
| **Man $\alpha$(1-6) Man (1-6)** | $\phi$ | $\psi$ | $\omega$ | **Pop (%)** |
| Cluster 1 | 71.9 (10.2) | -166.8 (16.7) | 55.1 (10.1) | 68.2 |
| Cluster 2 | 69.2 (8.2) | -173.8 (12.6) | - 83.8 (11.2) | 4.0 |
| Cluster 3 | 68.9 (7.6) | -120.6 (11.9) | - 65.4 (10.0) | 4.0 |
| Cluster 4 | 69.0 (8.9) | -175.7 (23.2) | -165.9 (10.6) | 4.5 |
| Cluster 5 | 78.2 (16.7) | 86.2 (10.1) | 49.8 (10.1) | 12.3 |
| Cluster 6 | 71.9 (10.4) | 82.6 (12.5) | -166.3 (17.7) | 6.1 |
| Cluster 7 | 84.9 (5.7) | - 78.2 (7.5) | - 79.57 (6.4) | 1.0 |
| **Man $\alpha$(1-2) Man (1-6)(1-6)** | $\phi$ | $\psi$ | $\omega$ | **Pop (%)** |
| Cluster 1 | 74.7 (8.6) | 152.1 (15.1) | - | 79.1 |
| Cluster 2 | 71.3 (9.6) | 106.4 (12.2) | - | 20.9 |

### 6 Man 7 I

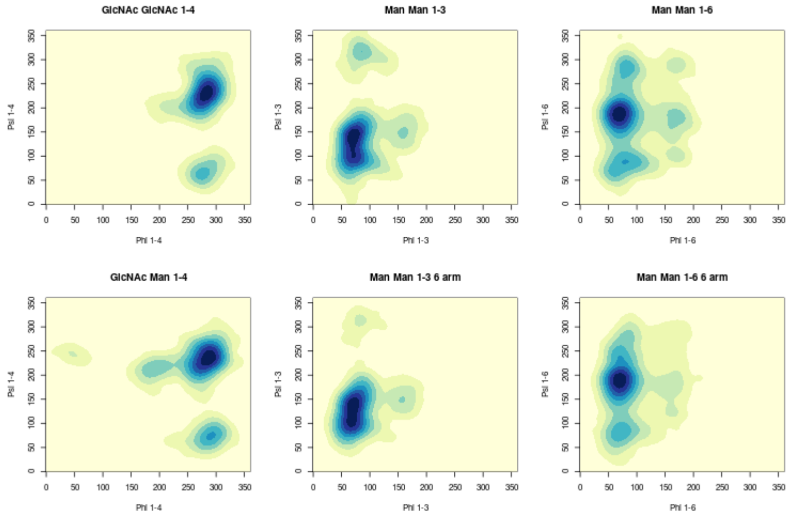


Figure 6: 2D Kernel density estimates for the $\phi$ / $\psi$ angle distributions for Man 7 I.

Table 5: The $\phi$ / $\psi$ / $\omega$ angle distributions for Man 7 I.

| **GlcNAc $\beta$(1-4) GlcNAc** | $\phi$ | $\psi$ | $\omega$ | **Pop (%)** |
|---|---|---|---|---|
| Cluster 1 | -77.9 (10.9) | -130.2 (17.5) | - | 95.2 |
| Cluster 2 | -82.1 (10.8) | 65.5 (11.4) | - | 4.8 |
| **Man $\beta$(1-4) GlcNAc** | $\phi$ | $\psi$ | $\omega$ | **Pop (%)** |
| Cluster 1 | -76.6 (12.8) | -125.5 (15.4) | - | 89.8 |
| Cluster 2 | -69.3 (10.9) | 73.5 (11.3) | - | 7.7 |
| Cluster 3 | -167.7 (12.2) | -145.2 (8.3) | - | 2.5 |
| **Man $\alpha$(1-3) Man (1-3)** | $\phi$ | $\psi$ | $\omega$ | **Pop (%)** |
| Cluster 1 | 72.1 (9.0) | 141.5 (14.1) | - | 63.3 |
| Cluster 2 | 71.3 (10.3) | 97.6 (10.7) | - | 36.7 |
| **Man $\alpha$(1-3) Man (1-6)** | $\phi$ | $\psi$ | $\omega$ | **Pop (%)** |
| Cluster 1 | 71.9 (9.2) | 138.6 (14.8) | - | 61.1 |
| Cluster 2 | 67.5 (9.9) | 99.5 (10.4) | - | 38.9 |
| **Man $\alpha$(1-6) Man** | $\phi$ | $\psi$ | $\omega$ | **Pop (%)** |
| Cluster 1 | 70.7 (10.8) | -172.3 (15.8) | 55.4 (11.2) | 46.2 |
| Cluster 2 | 68.0 (10.0) | -175.6 (14.1) | -175.6 (12.3) | 33.1 |
| Cluster 3 | 76.6 (16.5) | 84.9 (14.7) | 49.8 (9.8) | 11.7 |
| Cluster 4 | 82.0 (8.3) | -76.0 (10.9) | -148.8 (10.6) | 4.6 |
| Cluster 5 | 71.5 (9.1) | -178.9 (14.6) | -71.8 (11.5) | 4.4 |
| **Man $\alpha$(1-6) Man (1-6)** | $\phi$ | $\psi$ | $\omega$ | **Pop (%)** |
| Cluster 1 | 70.2 (10.4) | -171.5 (15.4) | 54.4 (10.4) | 78.1 |
| Cluster 2 | 69.1 (8.3) | -175.9 (12.8) | -79.3 (12.2) | 5.7 |
| Cluster 3 | 69.2 (7.8) | -118.2 (13.2) | -65.1 (10.1) | 5.4 |
| Cluster 4 | 69.8 (8.5) | -172.0 (18.5) | -165.6 (10.0) | 7.4 |
| Cluster 5 | 71.3 (9.5) | 84.0 (10.3) | 47.7 (8.0) | 3.4 |
| **Man $\alpha$(1-2) Man (1-3)** | $\phi$ | $\psi$ | $\omega$ | **Pop (%)** |
| Cluster 1 | 74.6 (9.0) | 152.5 (15.4) | - | 72.0 |
| Cluster 2 | 70.0 (9.7) | 105.3 (11.6) | - | 28.0 |
| **Man $\alpha$(1-2) Man (1-3)** | $\phi$ | $\psi$ | $\omega$ | **Pop (%)** |
| Cluster 1 | 74.1 (8.8) | 151.4 (15.0) | - | 76.0 |
| Cluster 2 | 70.0 (9.3) | 106.5 (11.6) | - | 24.0 |

### 7 Man 7 II

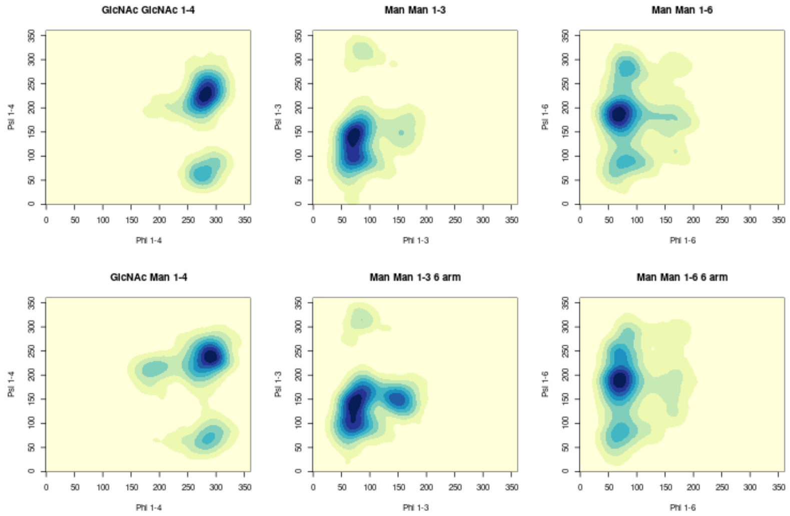


Figure 7: 2D Kernel density estimates for the $\phi$ / $\psi$ angle distributions for Man 7 II.

Table 6: The $\phi$ / $\psi$ / $\omega$ angle distributions for Man 7 II.

| **GlcNAc $\beta$(1-4) GlcNAc** | $\phi$ | $\psi$ | $\omega$ | **Pop (%)** |
|---|---|---|---|---|
| Cluster 1 | -78.8 (10.2) | -131.6 (16.0) | - | 92.8 |
| Cluster 2 | -81.8 (11.8) | 65.5 (11.4) | - | 7.2 |
| **Man $\beta$(1-4) GlcNAc** | $\phi$ | $\psi$ | $\omega$ | **Pop (%)** |
| Cluster 1 | -71.7 (12.8) | -125.5 (15.4) | - | 92.3 |
| Cluster 2 | -72.5 (11.4) | 69.3 (11.6) | - | 5.0 |
| Cluster 3 | -170.2 (12.3) | -146.7 (8.0) | - | 2.7 |
| **Man $\alpha$(1-3) Man (1-3)** | $\phi$ | $\psi$ | $\omega$ | **Pop (%)** |
| Cluster 1 | 72.5 (9.2) | 142.0 (14.1) | - | 67.6 |
| Cluster 2 | 70.8 (10.2) | 98.2 (10.5) | - | 32.4 |
| **Man $\alpha$(1-3) Man (1-6)** | $\phi$ | $\psi$ | $\omega$ | **Pop (%)** |
| Cluster 1 | 75.9 (11.1) | 138.6 (14.8) | - | 59.0 |
| Cluster 2 | 70.4 (10.7) | 100.2 (9.6) | - | 26.5 |
| Cluster 3 | 148.9 (10.2) | 150.6 (9.7) | - | 14.5 |
| **Man $\alpha$(1-6) Man** | $\phi$ | $\psi$ | $\omega$ | **Pop (%)** |
| Cluster 1 | 71.1 (12.5) | -171.9 (15.6) | 56.1 (11.9) | 55.5 |
| Cluster 2 | 66.7 (10.2) | -177.4 (13.0) | -175.7 (12.4) | 31.7 |
| Cluster 3 | 79.6 (15.6) | 87.9 (13.3) | 50.1 (9.7) | 6.5 |
| Cluster 4 | 82.4 (8.8) | - 77.6 (10.8) | -150.0 (10.3) | 5.2 |
| Cluster 5 | 71.6 (7.2) | -177.4 (9.8) | - 71.7 (7.9) | 1.2 |
| **Man $\alpha$(1-6) Man (1-6)** | $\phi$ | $\psi$ | $\omega$ | **Pop (%)** |
| Cluster 1 | 70.3 (10.5) | -170.8 (16.1) | 54.8 (10.5) | 71.9 |
| Cluster 2 | 69.8 (7.8) | -179.8 (14.0) | - 74.4 (9.6) | 5.0 |
| Cluster 3 | 70.0 (7.5) | -117.8 (13.4) | - 67.3 (10.1) | 8.5 |
| Cluster 4 | 70.7 (9.8) | -172.4 (24.5) | -163.9 (12.9) | 8.3 |
| Cluster 5 | 71.0 (12.2) | 84.4 (13.6) | 48.8 (9.9) | 4.4 |
| Cluster 6 | 67.8 (10.7) | 73.6 (12.5) | -177.1 (10.1) | 1.9 |
| **Man $\alpha$(1-2) Man (1-3)** | $\phi$ | $\psi$ | $\omega$ | **Pop (%)** |
| Cluster 1 | 74.2 (8.7) | 150.6 (14.7) | - | 72.5 |
| Cluster 2 | 70.0 (9.2) | 107.4 (11.6) | - | 27.5 |
| **Man $\alpha$(1-2) Man (1-3)(1-3)** | $\phi$ | $\psi$ | $\omega$ | **Pop (%)** |
| Cluster 1 | 72.4 (8.1) | 149.5 (13.6) | - | 75.9 |
| Cluster 2 | 69.7 (8.2) | 111.7 (10.6) | - | 24.1 |

### 8 Man 7 III

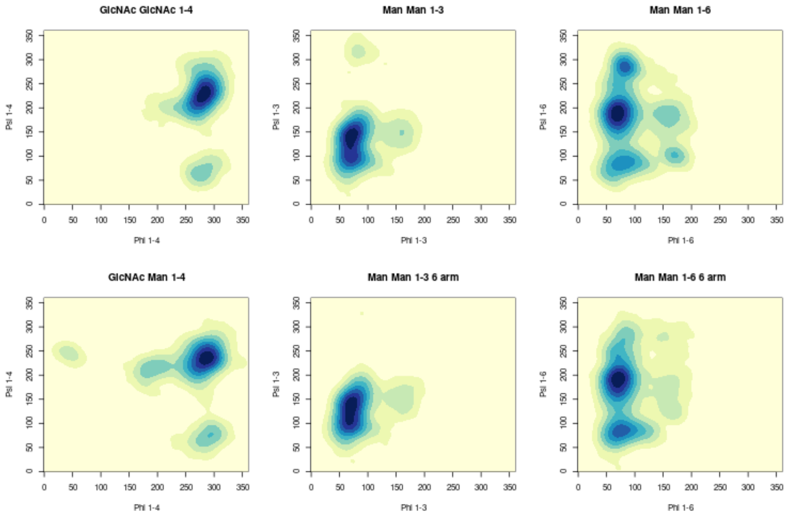


Figure 8: 2D Kernel density estimates for the $\phi$ / $\psi$ angle distributions for Man 7 III.

Table 7: The $\phi$ / $\psi$ / $\omega$ angle distributions for Man 7 III.

| **GlcNAc $\beta$(1-4) GlcNAc** | $\phi$ | $\psi$ | $\omega$ | **Pop (%)** |
|---|---|---|---|---|
| Cluster 1 | -78.9 (11.5) | -131.9 (18.3) | - | 97.7 |
| Cluster 2 | -82.7 (9.8) | 65.3 (9.0) | - | 2.3 |
| **Man $\beta$(1-4) GlcNAc** | $\phi$ | $\psi$ | $\omega$ | **Pop (%)** |
| Cluster 1 | -75.4 (12.7) | -125.3 (14.8) | - | 94.0 |
| Cluster 2 | -68.4 (10.4) | 73.5 (10.4) | - | 3.1 |
| Cluster 3 | -167.5 (12.4) | -145.6 (8.0) | - | 2.9 |
| **Man $\alpha$(1-3) Man (1-3)** | $\phi$ | $\psi$ | $\omega$ | **Pop (%)** |
| Cluster 1 | 72.1 (9.0) | 141.5 (14.5) | - | 66.5 |
| Cluster 2 | 70.4 (9.9) | 98.3 (10.4) | - | 33.5 |
| **Man $\alpha$(1-3) Man (1-6)** | $\phi$ | $\psi$ | $\omega$ | **Pop (%)** |
| Cluster 1 | 71.8 (9.1) | 139.1 (14.6) | - | 65.1 |
| Cluster 2 | 67.8 (9.5) | 100.4 (9.8) | - | 34.9 |
| **Man $\alpha$(1-6) Man** | $\phi$ | $\psi$ | $\omega$ | **Pop (%)** |
| Cluster 1 | 71.1 (10.7) | -172.4 (16.1) | 56.0 (10.7) | 42.6 |
| Cluster 2 | 67.2 (9.4) | -175.5 (15.4) | -176.2 (11.7) | 20.2 |
| Cluster 3 | 78.0 (16.2) | 85.3 (13.5) | 49.5 (10.3) | 14.5 |
| Cluster 4 | 82.4 (7.7) | - 74.3 (9.9) | -151.0 (9.4) | 11.3 |
| Cluster 5 | 73.3 (9.5) | -160.8 (20.5) | - 66.1 (12.0) | 7.8 |
| Cluster 6 | 72.8 (8.6) | - 98.6 (10.0) | - 73.0 (9.1) | 1.9 |
| Cluster 7 | 169.8 (7.8) | 102.0 (7.2) | 171.2 (7.6) | 1.7 |
| **Man $\alpha$(1-6) Man (1-6)** | $\phi$ | $\psi$ | $\omega$ | **Pop (%)** |
| Cluster 1 | 71.3 (10.3) | -169.3 (16.5) | 55.0 (10.3) | 56.7 |
| Cluster 2 | 69.2 (9.1) | -174.3 (15.0) | - 79.7 (13.6) | 6.2 |
| Cluster 3 | 71.2 (8.9) | -113.5 (17.1) | - 67.5 (11.4) | 5.0 |
| Cluster 4 | 69.4 (9.6) | -178.5 (21.7) | -166.2 (12.1) | 9.2 |
| Cluster 5 | 79.6 (17.5) | 85.8 (11.4) | 49.8 (9.6) | 14.5 |
| Cluster 6 | 71.9 (11.1) | 80.9 (11.2) | -168.0 (14.6) | 8.4 |
| **Man $\alpha$(1-2) Man (1-3)** | $\phi$ | $\psi$ | $\omega$ | **Pop (%)** |
| Cluster 1 | 74.2 (8.7) | 150.4 (14.8) | - | 72.1 |
| Cluster 2 | 69.8 (9.1) | 107.5 (11.3) | - | 27.9 |
| **Man $\alpha$(1-2) Man (1-6)(1-6)** | $\phi$ | $\psi$ | $\omega$ | **Pop (%)** |
| Cluster 1 | 74.5 (8.5) | 151.5 (14.5) | - | 78.0 |
| Cluster 2 | 70.9 (9.1) | 106.7 (11.8) | - | 22.0 |

### 9 Man 7 IV

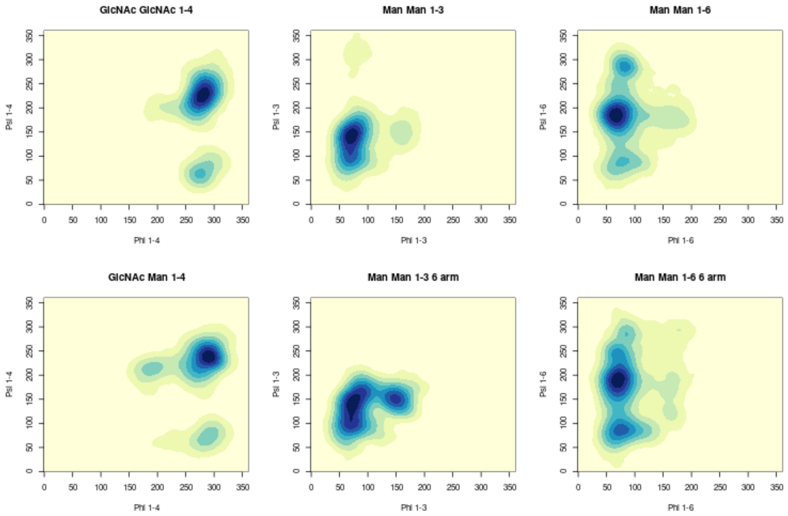


Figure 9: 2D Kernel density estimates for the $\phi$ / $\psi$ angle distributions for Man 7 IV.

Table 8: The $\phi$ / $\psi$ / $\omega$ angle distributions for Man 7 IV.

| **GlcNAc $\beta$(1-4) GlcNAc** | $\phi$ | $\psi$ | $\omega$ | **Pop (%)** |
|---|---|---|---|---|
| Cluster 1 | -79.8 (10.2) | -133.1 (16.5) | - | 95.8 |
| Cluster 2 | -82.3 (10.6) | 65.8 (11.8) | - | 4.2 |
| **Man $\beta$(1-4) GlcNAc** | $\phi$ | $\psi$ | $\omega$ | **Pop (%)** |
| Cluster 1 | -71.8 (12.0) | -122.9 (14.1) | - | 95.3 |
| Cluster 2 | -70.0 (10.7) | 72.3 (10.9) | - | 2.6 |
| Cluster 3 | -169.2 (10.7) | -145.3 (7.7 | - | 2.1 |
| **Man $\alpha$(1-3) Man (1-3)** | $\phi$ | $\psi$ | $\omega$ | **Pop (%)** |
| Cluster 1 | 72.9 (9.1) | 142.4 (14.1) | - | 77.3 |
| Cluster 2 | 69.3 (9.1) | 100.1 (10.3) | - | 22.7 |
| **Man $\alpha$(1-3) Man (1-6)** | $\phi$ | $\psi$ | $\omega$ | **Pop (%)** |
| Cluster 1 | 76.6 (11.8) | 145.7 (16.9) | - | 56.9 |
| Cluster 2 | 70.6 (10.7) | 100.0 (9.7) | - | 23.7 |
| Cluster 3 | 148.4 (10.7) | 151.5 (10.3) | - | 19.4 |
| **Man $\alpha$(1-6) Man** | $\phi$ | $\psi$ | $\omega$ | **Pop (%)** |
| Cluster 1 | 70.5 (12.4) | -172.4 (15.5) | 55.7 (11.5) | 46.0 |
| Cluster 2 | 66.3 (9.9) | -178.1 (13.4) | -176.7 (11.7) | 40.8 |
| Cluster 3 | 78.4 (14.7) | 87.6 (12.9) | 50.3 (9.7) | 5.1 |
| Cluster 4 | 82.0 (7.5) | - 74.7 (10.1) | -151.1 (9.8) | 8.1 |
| **Man $\alpha$(1-6) Man (1-6)** | $\phi$ | $\psi$ | $\omega$ | **Pop (%)** |
| Cluster 1 | 71.1 (10.3) | -170.2 (16.6) | 54.9 (10.4) | 61.8 |
| Cluster 2 | 70.2 (7.3) | 177.6 (10.8) | - 72.6 (8.8) | 3.0 |
| Cluster 3 | 70.1 (7.5) | -118.6 (12.0) | - 66.9 (9.8) | 7.5 |
| Cluster 4 | 69.9 (9.2) | -179.5 (22.1) | -165.8 (11.2) | 5.4 |
| Cluster 5 | 79.8 (16.5) | 86.1 (11.9) | 50.5 (9.5) | 12.5 |
| Cluster 6 | 71.6 (10.2) | 85.4 (12.6) | -158.7 (21.5) | 8.8 |
| Cluster 7 | 84.9 (5.7) | - 76.2 (7.5) | - 80.6 (6.1) | 1.2 |
| **Man $\alpha$(1-2) Man (1-3)** | $\phi$ | $\psi$ | $\omega$ | **Pop (%)** |
| Cluster 1 | 74.5 (8.8) | 151.5 (14.9) | - | 77.2 |
| Cluster 2 | 70.6 (9.3) | 106.2 (12.2) | - | 22.8 |
| **Man $\alpha$(1-2) Man (1-6)(1-6)** | $\phi$ | $\psi$ | $\omega$ | **Pop (%)** |
| Cluster 1 | 72.2 (8.0) | 149.4 (13.6) | - | 74.1 |
| Cluster 2 | 69.7 (8.1) | 112.1 (10.4) | - | 25.9 |

### 10 Man 8 I

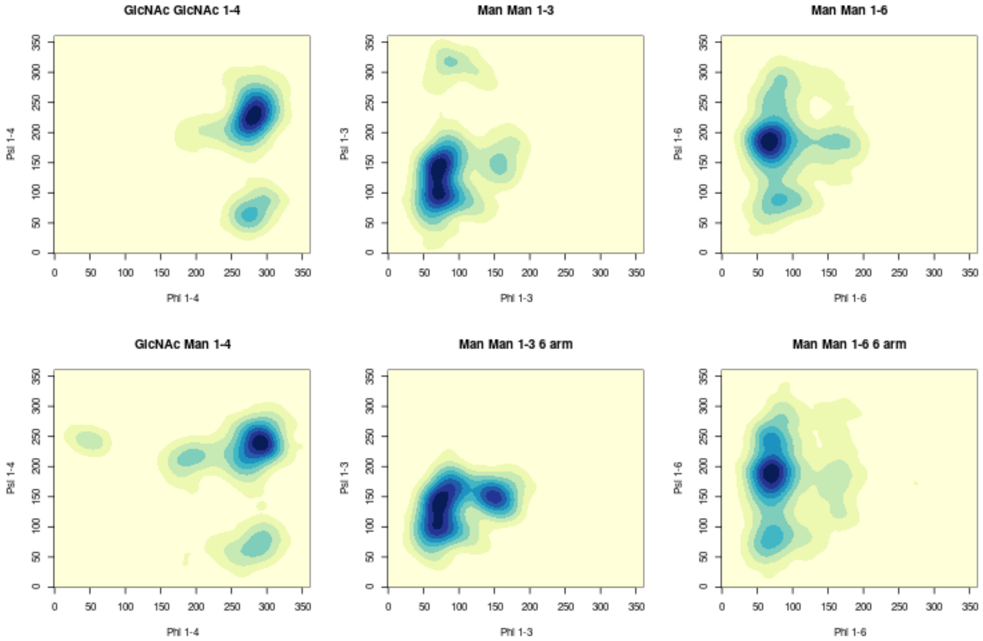


Figure 10: 2D Kernel density estimates for the $\phi$ / $\psi$ angle distributions for Man 8 I.

Table 9: The $\phi$ / $\psi$ / $\omega$ angle distributions for Man 8 I

| **GlcNAc $\beta$(1-4) GlcNAc** | $\phi$ | $\psi$ | $\omega$ | **Pop (%)** |
|---|---|---|---|---|
| Cluster 1 | -78.8 (10.1) | -131.7 (16.2) | - | 94.9 |
| Cluster 2 | -82.2 (11.5) | 66.7 (12.6) | - | 5.1 |
| **Man $\beta$(1-4) GlcNAc** | $\phi$ | $\psi$ | $\omega$ | **Pop (%)** |
| Cluster 1 | -71.3 (11.7) | -122.2 (14.1) | - | 94.8 |
| Cluster 2 | -72.4 (10.9) | 70.3 (11.3) | - | 3.2 |
| Cluster 3 | -167.7 (11.9) | -145.0 (8.1) | - | 2.2 |
| **Man $\alpha$(1-3) Man (1-3)** | $\phi$ | $\psi$ | $\omega$ | **Pop (%)** |
| Cluster 1 | 72.2 (9.0) | 141.3 (14.6) | - | 58.5 |
| Cluster 2 | 71.2 (10.4) | 96.9 (10.9) | - | 41.5 |
| **Man $\alpha$(1-3) Man (1-6)** | $\phi$ | $\psi$ | $\omega$ | **Pop (%)** |
| Cluster 1 | 75.8 (11.6) | 143.9 (16.7) | - | 53.5 |
| Cluster 2 | 70.2 (10.7) | 99.9 (9.5) | - | 26.5 |
| Cluster 2 | 70.2 (10.7) | 150.2 (10.0) | - | 19.9 |
| **Man $\alpha$(1-6) Man** | $\phi$ | $\psi$ | $\omega$ | **Pop (%)** |
| Cluster 1 | 69.5 (12.3) | -173.0 (15.5) | 54.1 (12.6) | 55.3 |
| Cluster 2 | 66.3 (9.8) | -178.0 (12.6) | -176.1 (12.1) | 38.2 |
| Cluster 3 | 81.4 (13.1) | 88.3 (11.3) | 49.3 (10.3) | 4.8 |
| Cluster 4 | 82.0 (8.3) | -76.0 (10.9) | -148.8 (10.6) | 0.0 |
| Cluster 5 | 75.0 (7.31) | -171.1 (9.5) | -64.8 (8.2) | 1.7 |
| **Man $\alpha$(1-6) Man (1-6)** | $\phi$ | $\psi$ | $\omega$ | **Pop (%)** |
| Cluster 1 | 70.3 (10.4) | -171.5 (15.8) | 54.7 (10.7) | 74.9 |
| Cluster 2 | 71.2 (7.4) | -178.9 (14.1) | -73.2 (8.8) | 5.4 |
| Cluster 3 | 69.4 (7.2) | -117.9 (12.3) | -67.0 (9.7) | 9.4 |
| Cluster 4 | 69.8 (9.2) | -175.0 (19.6) | -164.4 (11.3) | 6.2 |
| Cluster 5 | 70.3 (10.6) | 83.9 (12.5) | 48.0 (9.3) | 4.1 |
| **Man $\alpha$(1-2) Man (1-3)** | $\phi$ | $\psi$ | $\omega$ | **Pop (%)** |
| Cluster 1 | 75.4 (9.0) | 151.7 (15.6) | - | 70.8 |
| Cluster 2 | 71.1 (9.7) | 105.9 (11.6) | - | 29.2 |
| **Man $\alpha$(1-2) Man (1-3)** | $\phi$ | $\psi$ | $\omega$ | **Pop (%)** |
| Cluster 1 | 73.9 (8.8) | 151.2 (15.2) | - | 74.8 |
| Cluster 2 | 70.1 (9.2) | 106.6 (11.7) | - | 25.2 |
| **Man $\alpha$(1-2) Man (1-6)** | $\phi$ | $\psi$ | $\omega$ | **Pop (%)** |
| Cluster 1 | 73.9 (8.8) | 151.2 (15.2) | - | 74.8 |
| Cluster 2 | 70.1 (9.2) | 106.6 (11.7) | - | 25.2 |

### 11 Man 8 II

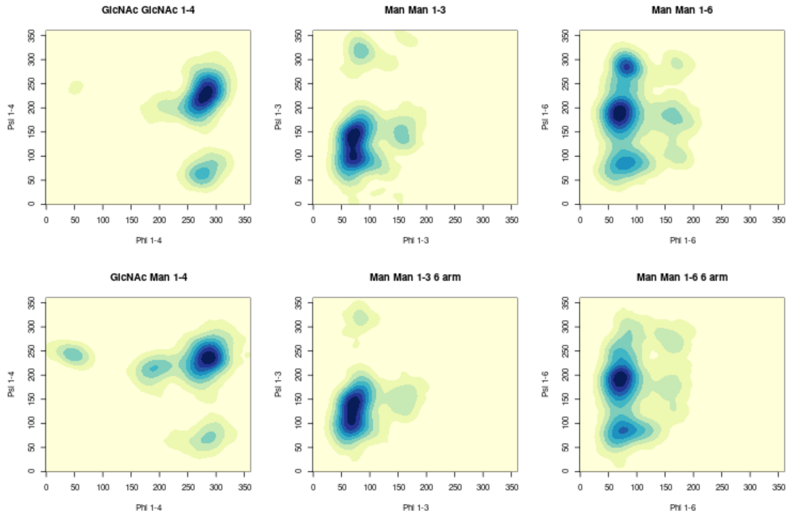


Figure 11: 2D Kernel density estimates for the $\phi$ / $\psi$ angle distributions for Man 8 II.

Table 10: The $\phi$ / $\psi$ / $\omega$ angle distributions for Man 8 II.

| **GlcNAc $\beta$(1-4) GlcNAc** | $\phi$ | $\psi$ | $\omega$ | **Pop (%)** |
|---|---|---|---|---|
| Cluster 1 | -79.4 (11.4) | -132.7 (17.5) | - | 95.4 |
| Cluster 2 | -82.4 (11.2) | 65.6 (11.0) | - | 4.6 |
| **Man $\beta$(1-4) GlcNAc** | $\phi$ | $\psi$ | $\omega$ | **Pop (%)** |
| Cluster 1 | -75.1 (12.4) | -125.0 (14.9) | - | 93.9 |
| Cluster 2 | -72.8 (8.2) | 69.6 (8.0) | - | 1.1 |
| Cluster 3 | -167.9 (12.5) | -145.3 (8.2) | - | 3.3 |
| Cluster 4 | 47.9 (8.5) | -117.23 (5.7) | - | 1.7 |
| **Man $\alpha$(1-3) Man (1-3)** | $\phi$ | $\psi$ | $\omega$ | **Pop (%)** |
| Cluster 1 | 72.2 (9.1) | 141.7 (14.6) | - | 61.0 |
| Cluster 2 | 71.0 (10.4) | 97.0 (10.8) | - | 39.0 |
| **Man $\alpha$(1-3) Man (1-6)** | $\phi$ | $\psi$ | $\omega$ | **Pop (%)** |
| Cluster 1 | 72.1 (9.2) | 139.7 (14.8) | - | 64.7 |
| Cluster 2 | 67.4 (9.6) | 100.2 (10.3) | - | 35.3 |
| **Man $\alpha$(1-6) Man** | $\phi$ | $\psi$ | $\omega$ | **Pop (%)** |
| Cluster 1 | 71.1 (10.8) | -171.6 (16.3) | 55.4 (11.2) | 47.5 |
| Cluster 2 | 67.4 (9.9) | -173.9 (14.9) | -175.6 (12.0) | 18.0 |
| Cluster 3 | 79.6 (15.1) | 85.7 (13.3) | 48.9 (10.4) | 11.8 |
| Cluster 4 | 81.9 (7.6) | - 74.7 (9.6) | -150.7 (9.4) | 16.6 |
| Cluster 5 | 72.6 (8.9) | -164.8 (19.3) | - 67.6 (11.7) | 6.0 |
| **Man $\alpha$(1-6) Man (1-6)** | $\phi$ | $\psi$ | $\omega$ | **Pop (%)** |
| Cluster 1 | 71.4 (10.2) | -168.5 (16.4) | 54.8 (10.3) | 67.5 |
| Cluster 2 | 69.4 (8.8) | -175.8 (13.9) | - 80.6 (13.5) | 5.0 |
| Cluster 3 | 69.1 (6.3) | -119.2 (8.8) | - 63.6 (7.6) | 2.4 |
| Cluster 4 | 69.5 (9.2) | -175.5 (21.3) | -166.7 (10.9) | 6.7 |
| Cluster 5 | 79.0 (16.6) | 86.2 (11.7) | 50.1 (9.8) | 13.1 |
| Cluster 6 | 71.6 (10.6) | 81.9 (11.4) | -166.1 (13.5) | 5.4 |
| **Man $\alpha$(1-2) Man (1-3)** | $\phi$ | $\psi$ | $\omega$ | **Pop (%)** |
| Cluster 1 | 74.7 (9.0) | 152.3 (15.5) | - | 62.9 |
| Cluster 2 | 72.2 (10.0) | 97.8 (17.8) | - | 37.1 |
| **Man $\alpha$(1-2) Man (1-3)** | $\phi$ | $\psi$ | $\omega$ | **Pop (%)** |
| Cluster 1 | 74.3 (8.4) | 152.1 (15.1) | - | 75.6 |
| Cluster 2 | 70.0 (9.2) | 106.6 (11.7) | - | 24.4 |
| **Man $\alpha$(1-2) Man (1-6)(1-6)** | $\phi$ | $\psi$ | $\omega$ | **Pop (%)** |
| Cluster 1 | 74.3 (8.8) | 152.1 (15.2) | - | 75.6 |
| Cluster 2 | 70.0 (9.2) | 106.7 (11.7) | - | 24.4 |

### 12 Man 8 III

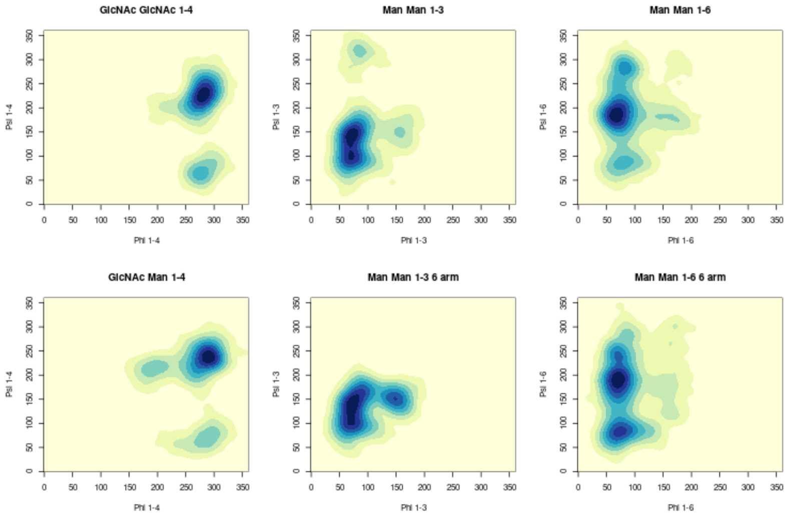


Figure 12: 2D Kernel density estimates for the $\phi$ / $\psi$ angle distributions for Man 8 III.

Table 11: The $\phi$ / $\psi$ / $\omega$ angle distributions for Man 8 III.

| **GlcNAc $\beta$(1-4) GlcNAc** | $\phi$ | $\psi$ | $\omega$ | **Pop (%)** |
|---|---|---|---|---|
| Cluster 1 | -79.3 (10.5) | -132.7 (16.4) | - | 94.0 |
| Cluster 2 | -82.3 (10.6) | 65.9 (11.1) | - | 6.0 |
| **Man $\beta$(1-4) GlcNAc** | $\phi$ | $\psi$ | $\omega$ | **Pop (%)** |
| Cluster 1 | -71.8 (11.9) | -123.4 (13.9) | - | 94.0 |
| Cluster 2 | -71.6 (10.3) | 69.8 (10.5) | - | 3.2 |
| Cluster 3 | -167.7 (11.2) | -145.4 (7.7) | - | 2.7 |
| **Man $\alpha$(1-3) Man (1-3)** | $\phi$ | $\psi$ | $\omega$ | **Pop (%)** |
| Cluster 1 | 72.4 (9.0) | 142.1 (14.1) | - | 61.2 |
| Cluster 2 | 71.8 (9.8) | 97.3 (10.2) | - | 37.8 |
| **Man $\alpha$(1-3) Man (1-6)** | $\phi$ | $\psi$ | $\omega$ | **Pop (%)** |
| Cluster 1 | 75.4 (11.1) | 143.5 (16.2) | - | 58.3 |
| Cluster 2 | 71.1 (10.2) | 100.6 (9.1) | - | 27.7 |
| Cluster 3 | 148.3 (9.3) | 151.5 (8.7) | - | 14.1 |
| **Man $\alpha$(1-6) Man** | $\phi$ | $\psi$ | $\omega$ | **Pop (%)** |
| Cluster 1 | 70.6 (11.7) | -172.7 (14.8) | 56.2 (11.7) | 36.7 |
| Cluster 2 | 66.3 (9.8) | -178.4 (12.7) | -176.7 (11.7) | 35.9 |
| Cluster 3 | 80.2 (11.7) | 86.2 (9.7) | 48.0 (9.3) | 4.3 |
| Cluster 4 | 82.9 (8.0) | - 76.2 (9.39) | -150.8 (9.31) | 8.1 |
| Cluster 5 | 74.5 (9.7) | -152.8 (16.7) | - 64.6 (11.8) | 13.0 |
| Cluster 6 | 72.4 (7.9) | -100.0 (8.2) | - 72.1 (8.0) | 2.2 |
| **Man $\alpha$(1-6) Man (1-6)** | $\phi$ | $\psi$ | $\omega$ | **Pop (%)** |
| Cluster 1 | 71.2 (10.1) | -170.4 (16.0) | 55.1 (10.3) | 50.0 |
| Cluster 2 | 70.2 (8.5) | -178.3 (14.4) | - 76.2 (10.7) | 5.9 |
| Cluster 3 | 69.9 (6.8) | -117.7 (10.2) | - 66.0 (8.5) | 8.2 |
| Cluster 4 | 69.5 (8.4) | 179.8 (16.4) | -166.4 (9.6) | 5.9 |
| Cluster 5 | 79.4 (15.8) | 85.4 (11.0) | 49.5 (9.4) | 12.0 |
| Cluster 6 | 71.7 (11.3) | 81.4 (11.8) | -166.5 (14.2) | 18.0 |
| **Man $\alpha$(1-2) Man (1-3)** | $\phi$ | $\psi$ | $\omega$ | **Pop (%)** |
| Cluster 1 | 73.7 (8.6) | 149.5 (14.3) | - | 70.4 |
| Cluster 2 | 69.7( 8.9) | 107.5 (11.3) | - | 29.5 |
| **Man $\alpha$(1-2) Man (1-6)(1-3)** | $\phi$ | $\psi$ | $\omega$ | **Pop (%)** |
| Cluster 1 | 72.3 (7.9) | 149.5 (13.3) | - | 72.2 |
| Cluster 2 | 69.9 (8.1) | 111.1 (10.3) | - | 27.8 |
| **Man $\alpha$(1-2) Man (1-6)(1-6)** | $\phi$ | $\psi$ | $\omega$ | **Pop (%)** |
| Cluster 1 | 72.3 (7.9) | 149.5 (13.3) | - | 72.2 |
| Cluster 2 | 69.1 (8.1) | 111.0 (10.4) | - | 27.8 |

### 13 Man 9

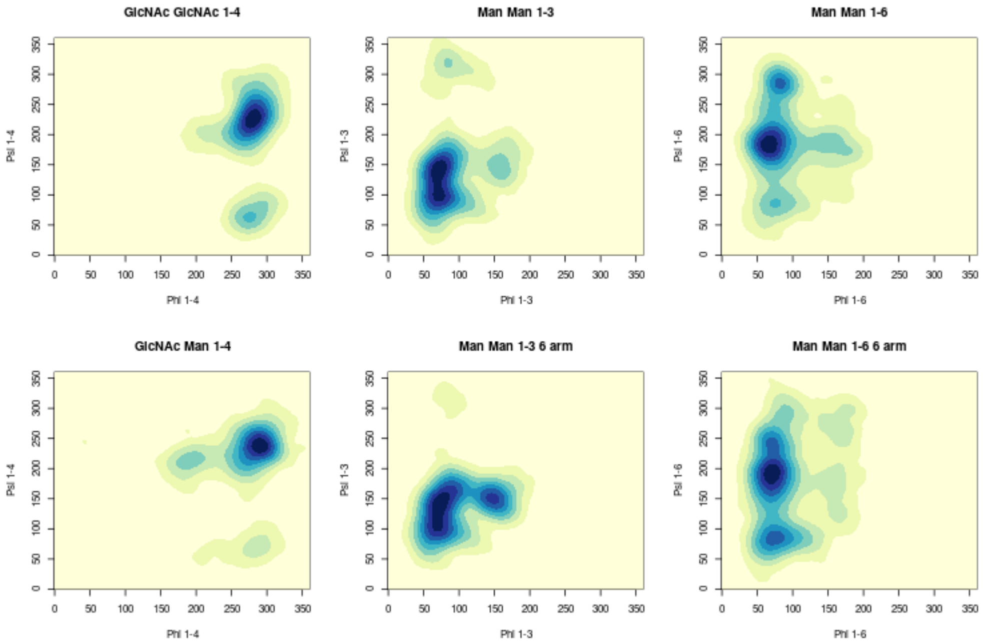


Figure 13: 2D Kernel density estimates for the $\phi$ / $\psi$ angle distributions for Man 8.

Table 12: The $\phi$ / $\psi$ / $\omega$ angle distributions for Man 9

| **GlcNAc $\beta$(1-4) GlcNAc** | $\phi$ | $\psi$ | $\omega$ | **Pop (%)** |
|---|---|---|---|---|
| Cluster 1 | -78.8 (10.5) | -131.7 (17.8) | - | 95.8 |
| Cluster 2 | -82.3 (11.8) | 65.9 (11.5) | - | 4.2 |
| **Man $\beta$(1-4) GlcNAc** | $\phi$ | $\psi$ | $\omega$ | **Pop (%)** |
| Cluster 1 | -71.8 (11.7) | -123.1 (13.5) | - | 97.5 |
| Cluster 3 | -168.4 (12.5) | -146.1 (8.7) | - | 2.5 |
| **Man $\alpha$(1-3) Man (1-3)** | $\phi$ | $\psi$ | $\omega$ | **Pop (%)** |
| Cluster 1 | 72.3 (9.2) | 141.5 (14.7) | - | 58.5 |
| Cluster 2 | 71.4 (10.8) | 96.7 (11.1) | - | 41.5 |
| **Man $\alpha$(1-3) Man (1-6)** | $\phi$ | $\psi$ | $\omega$ | **Pop (%)** |
| Cluster 1 | 76.1 (12.1) | 144.8 (16.9) | - | 55.4 |
| Cluster 2 | 69.8 (10.9) | 100.2 (9.7) | - | 26.4 |
| Cluster 2 | 147.8 (10.9) | 150.8 (10.4) | - | 18.2 |
| **Man $\alpha$(1-6) Man** | $\phi$ | $\psi$ | $\omega$ | **Pop (%)** |
| Cluster 1 | 70.3 (12.6) | -173.3 (15.5) | 55.1 (12.4) | 38.0 |
| Cluster 2 | 66.0 (9.8) | -178.6 (13.0) | -176.9 (11.6) | 37.2 |
| Cluster 3 | 77.9 (11.2) | 86.9 (10.5) | 46.2 (9.3) | 2.8 |
| Cluster 4 | 82.0 (7.5) | -74.4 (9.6) | -151.1 (9.6) | 11.9 |
| Cluster 5 | 73.2 (9.4) | -157.3 (20.6) | -65.4 (11.3) | 7.0 |
| Cluster 6 | 75.9 (8.6) | -95.6 (9.7) | -74.8 (8.9) | 1.8 |
| Cluster 7 | 147.3 (9.3) | -171.6 (7.8) | -151.1 (7.4) | 1.3 |
| **Man $\alpha$(1-6) Man (1-6)** | $\phi$ | $\psi$ | $\omega$ | **Pop (%)** |
| Cluster 1 | 71.4 (10.2) | -168.8 (16.6) | 57.6 (10.3) | 56.9 |
| Cluster 2 | 71.2 (6.7) | -180.0 (14.6) | -73.2 (7.8) | 3.8 |
| Cluster 3 | 70.2 (7.4) | -118.1 (11.7) | -67.4 (10.0) | 11.6 |
| Cluster 4 | 69.2 (9.5) | -179.3 (22.9) | -165.7 (11.4) | 7.5 |
| Cluster 5 | 78.8 (15.92) | 86.3 (11.8) | 50.8 (9.65) | 4.1 |
| Cluster 6 | 72.6 (11.8) | 82.9 (11.8) | -163.9 (17.6) | 10.0 |
| **Man $\alpha$(1-2) Man (1-3)** | $\phi$ | $\psi$ | $\omega$ | **Pop (%)** |
| Cluster 1 | 75.4 (9.2) | 151.8 (15.7) | - | 69.6 |
| Cluster 2 | 71.1 (9.1) | 105.9 (11.5) | - | 30.4 |
| **Man $\alpha$(1-2) Man (1-3)** | $\phi$ | $\psi$ | $\omega$ | **Pop (%)** |
| Cluster 1 | 74.1 (8.8) | 151.7 (15.4) | - | 74.7 |
| Cluster 2 | 70.1 (9.6) | 106.3 (11.8) | - | 25.3 |
| **Man $\alpha$(1-2) Man (1-6)** | $\phi$ | $\psi$ | $\omega$ | **Pop (%)** |
| Cluster 1 | 74.1 (8.8) | 151.8 (15.4) | - | 74.7 |
| Cluster 2 | 70.1 (9.6) | 106.3 (11.7) | - | 25.3 |
| **Man $\alpha$(1-2) Man (1-6)(1-6)** | $\phi$ | $\psi$ | $\omega$ | **Pop (%)** |
| Cluster 1 | 74.4 (8.8) | 151.2 (15.1) | - | 76.5 |
| Cluster 2 | 70.5 (9.5) | 106.6 (12.2) | - | 23.5 |

### 14 Fc$\gamma$RC: Man5 N45

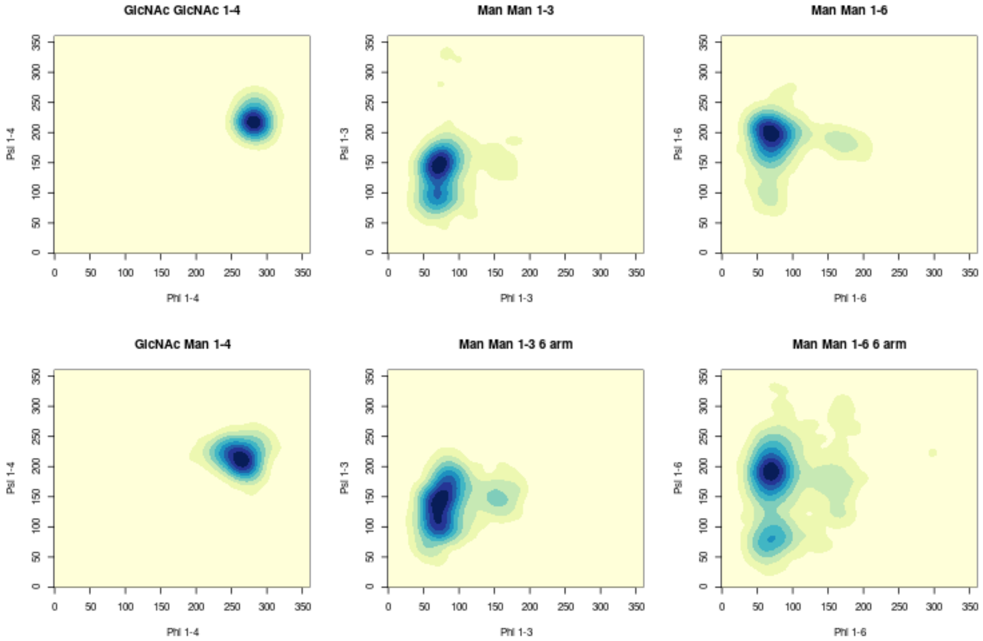


Figure 14: 2D Kernel density estimates for the $\phi$ / $\psi$ angle distributions for Man 5 (N45).

Table 13: The $\phi$ / $\psi$ / $\omega$ angle distributions for Man 5 on Fc$\gamma$-RC N45

| **GlcNAc $\beta$(1-4) GlcNAc** | $\phi$ | $\psi$ | $\omega$ | **Pop (%)** |
|---|---|---|---|---|
| Cluster 1 | -78.62 (6.6) | -142.4 (9.5) | - | 100 |
| **Man $\beta$(1-4) GlcNAc** | $\phi$ | $\psi$ | $\omega$ | **Pop (%)** |
| Cluster 1 | -96.9 (10.7) | -146.3 (11.3) | - | 93.0 |
| **Man $\alpha$(1-3) Man (1-3)** | $\phi$ | $\psi$ | $\omega$ | **Pop (%)** |
| Cluster 1 | 72.1 (8.1) | 145.9 (12.7) | - | 82.9 |
| Cluster 2 | 68.2 (8.4) | 97.2 (10.1) | - | 17.1 |
| **Man $\alpha$(1-3) Man (1-6)** | $\phi$ | $\psi$ | $\omega$ | **Pop (%)** |
| Cluster 1 | 75.4 (9.5) | 147.4 (15.0) | - | 65.9 |
| Cluster 2 | 70.6 (8.3) | 108.72 (10.6) | - | 34.1 |
| **Man $\alpha$(1-6) Man** | $\phi$ | $\psi$ | $\omega$ | **Pop (%)** |
| Cluster 1 | 69.1 (9.5) | -162.1 (13.9) | -172.2 (10.6) | 100 |
| **Man $\alpha$(1-6) Man (1-6)** | $\phi$ | $\psi$ | $\omega$ | **Pop (%)** |
| Cluster 1 | 69.7 (10.2) | -172.2 (15.1) | 53.8 (10.7) | 44.2 |
| Cluster 2 | 69.2 (9.4) | -166.3 (15.3) | -87.0 (15.8) | 36.9 |
| Cluster 3 | 70.6 (10.3) | -155.6 (18.9) | -161.1 (10.6) | 12.3 |
| Cluster 4 | 69.2 (10.6) | 79.9 (11.5) | -166.9 (12.5) | 6.7 |

### 15 Fc$\gamma$RC: Man5 N162

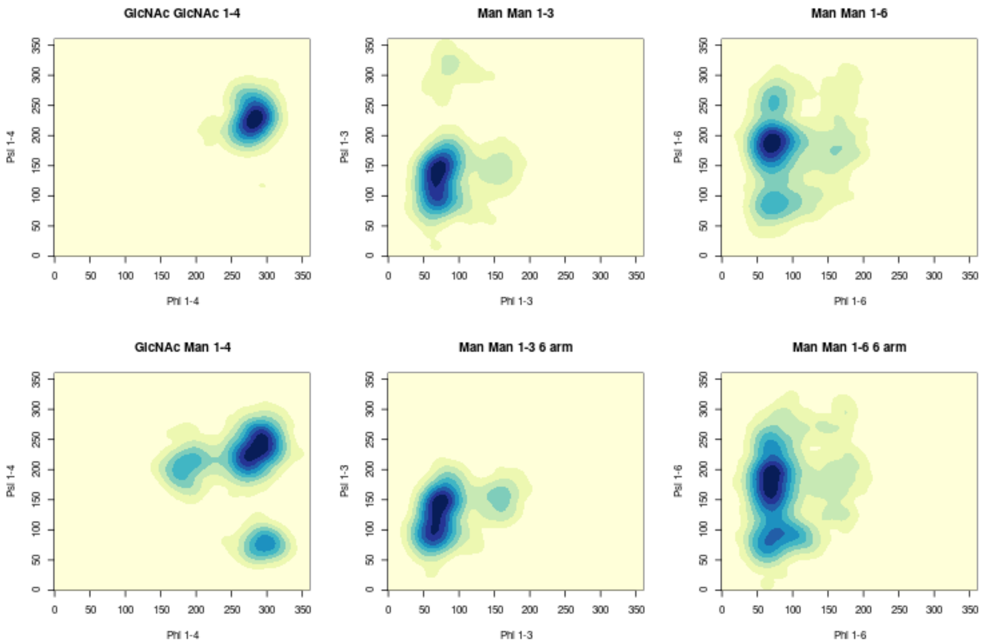


Figure 15: 2D Kernel density estimates for the $\phi$ / $\psi$ angle distributions for Man 5 on Fc$\gamma$-RC N162.

Table 14: The $\phi$ / $\psi$ / $\omega$ angle distributions for Man 5 on Fc$\gamma$-RC N162

| **Man $\alpha$(1-6) Man (1-6)** | $\phi$ | $\psi$ | $\omega$ | **Pop (%)** |
|---|---|---|---|---|
| Cluster 1 | -78.2 (9.3) | -132.8 (14.9) | - | 100 |
| **Man $\beta$(1-4) GlcNAc** | $\phi$ | $\psi$ | $\omega$ | **Pop (%)** |
| Cluster 1 | -74.12 (12.9) | -125.7 (17.2) | - | 86.7 |
| Cluster 2 | -62.9 (9.6) | 76.7 (8.9) | - | 8.3 |
| Cluster 3 | -175.3 (10.8) | -152.2 (11.8) | - | 5.0 |
| **Man $\alpha$(1-3) Man (1-3)** | $\phi$ | $\psi$ | $\omega$ | **Pop (%)** |
| Cluster 1 | 70.9 (8.8) | 141.5 (13.9) | - | 72.0 |
| Cluster 2 | 68.7 (8.9) | 100.9 (10.0) | - | 28.0 |
| **Man $\alpha$(1-3) Man (1-6)** | $\phi$ | $\psi$ | $\omega$ | **Pop (%)** |
| Cluster 1 | 73.5 (8.9) | 144.5 (11.1) | - | 52.1 |
| Cluster 2 | 65.9 (9.4) | 101.9 (13.6) | - | 47.9 |
| **Man $\alpha$(1-6) Man** | $\phi$ | $\psi$ | $\omega$ | **Pop (%)** |
| Cluster 1 | 69.1 (11.2) | -175.0 (16.1) | 52.3 (11.9) | 9.6 |
| Cluster 2 | 71.6 (11.9) | -172.8 (16.7) | -169.3 (14.9) | 82.5 |
| Cluster 3 | 74.0 (13.3) | 87.6,(12.8) | 52.6 (11.7) | 7.8 |
| **Man $\alpha$(1-6) Man (1-6)** | $\phi$ | $\psi$ | $\omega$ | **Pop (%)** |
| Cluster 1 | 70.3 (10.1) | 179.1 (22.7) | 57.4 (11.0) | 81.1 |
| Cluster 2 | 75.1 (15.8) | 89.4 (12.6) | 53.1 (9.7) | 18.9 |

### 16 Fc$\gamma$RC: Man9 N45

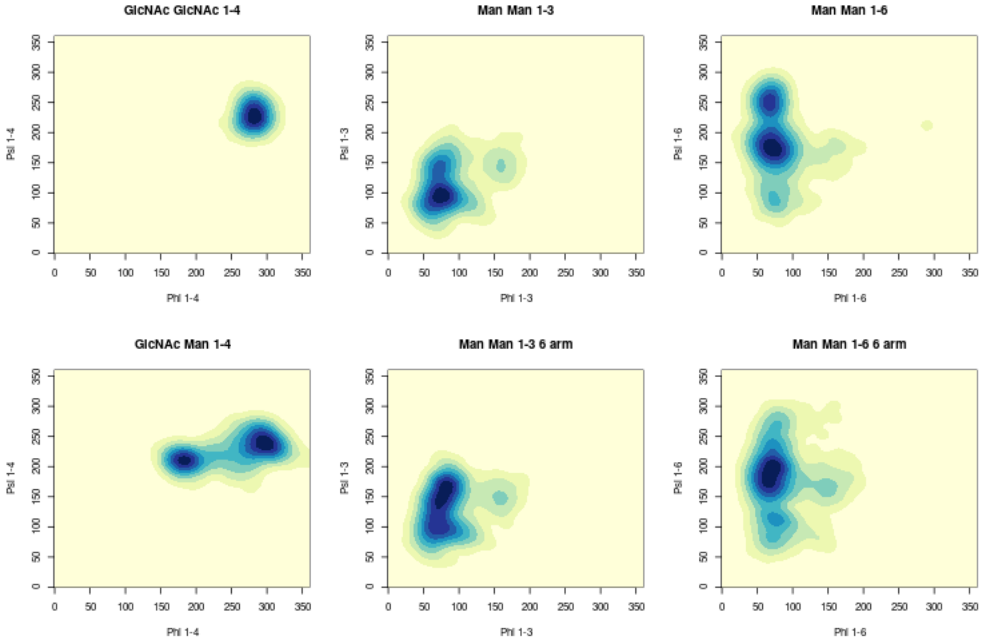


Figure 16: 2D Kernel density estimates for the $\phi$ / $\psi$ angle distributions for N45.

Table 15: The $\phi$ / $\psi$ / $\omega$ angle distributions for Man 9 on Fc$\gamma$-RC N45

| **GlcNAc $\beta$(1-4) GlcNAc** | $\phi$ | $\psi$ | $\omega$ | **Pop (%)** |
|---|---|---|---|---|
| Cluster 1 | -77.9 (7.5) | -132.6 (11.8) | - | 100 |
| **Man $\beta$(1-4) GlcNAc** | $\phi$ | $\psi$ | $\omega$ | **Pop (%)** |
| Cluster 1 | -65.9 (14.1) | -122.3 (11.6) | - | 64.6 |
| Cluster 2 | -176.2 (12.7) | -149.6 (7.1) | - | 35.4 |
| **Man $\alpha$(1-3) Man (1-3)** | $\phi$ | $\psi$ | $\omega$ | **Pop (%)** |
| Cluster 1 | 72.2 (7.9) | 138.1 (12.7) | - | 23.1 |
| Cluster 2 | 73.7 (11.4) | 93.4 (11.1) | - | 76.9 |
| **Man $\alpha$(1-3) Man (1-6)** | $\phi$ | $\psi$ | $\omega$ | **Pop (%)** |
| Cluster 1 | 78.3 (9.1) | 155.1 (15.3) | - | 60.2 |
| Cluster 2 | 70.8 (12.0) | 101.0 (12.4) | - | 39.8 |
| **Man $\alpha$(1-6) Man** | $\phi$ | $\psi$ | $\omega$ | **Pop (%)** |
| Cluster 1 | 71.2 (11.5) | 173.8 (13.1) | 59.4 (15.2) | 57.1 |
| Cluster 2 | 71.8 (8.7) | -173.6 (14.9) | -75.0 (11.3) | 15.1 |
| Cluster 3 | 67.1 (7.2) | -111.6,(12.1) | -69.1 (12.4) | 27.9 |
| **Man $\alpha$(1-6) Man (1-6)** | $\phi$ | $\psi$ | $\omega$ | **Pop (%)** |
| Cluster 1 | 71.0 (9.8) | 174.8 (14.8) | - 76.7 (14.1) | 17.2 |
| Cluster 2 | 65.7 (9.5) | -178.4 (22.2) | -170.2 (10.8) | 33.9 |
| Cluster 3 | 70.5 (10.2) | -164.1 (14.2) | 56.4 (11.7) | 42.9 |
| Cluster 4 | 76.1 (9.4) | 113.3 (8.3) | - 87.5 (7.9) | 5.9 |
| **Man $\alpha$(1-2) Man (1-3)** | $\phi$ | $\psi$ | $\omega$ | **Pop (%)** |
| Cluster 1 | 74.3 (8.4) | 148.2 (13.2) | - | 61.1 |
| Cluster 2 | 72.7 (10.1) | 105.3 (11.1) | - | 38.9 |
| **Man $\alpha$(1-2) Man (1-3)** | $\phi$ | $\psi$ | $\omega$ | **Pop (%)** |
| Cluster 1 | 73.2 (8.3) | 149.3 (14.3) | - | 69.1 |
| Cluster 2 | 70.1 (8.5) | 106.0 (10.9) | - | 30.9 |
| **Man $\alpha$(1-2) Man (1-6)** | $\phi$ | $\psi$ | $\omega$ | **Pop (%)** |
| Cluster 1 | 72.2 (7.6) | 146.9 (12.4) | - | 77.4 |
| Cluster 2 | 69.9 (7.9) | 107.1 (10.9) | - | 22.6 |
| **Man $\alpha$(1-2) Man (1-6)(1-6)** | $\phi$ | $\psi$ | $\omega$ | **Pop (%)** |
| Cluster 1 | 74.5 (9.0) | 151.9 (13.6) | - | 78.0 |
| Cluster 2 | 72.2 (8.1) | 104.2 (11.3) | - | 22.0 |

### 17 Fc$\gamma$RC: Man9 N162

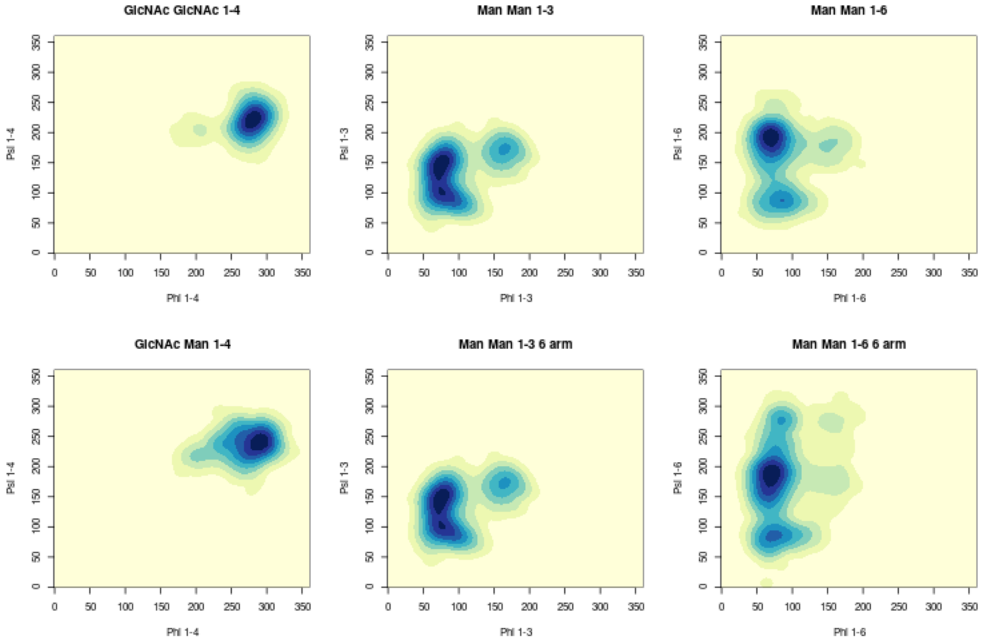


Figure 17: 2D Kernel density estimates for the $\phi$ / $\psi$ angle distributions for Man9 on the Fc$\gamma$-RC N162

Table 16: The $\phi$ / $\psi$ / $\omega$ angle distributions for Man 9 on Fc$\gamma$-RC N162

| **GlcNAc $\beta$(1-4) GlcNAc** | $\phi$ | $\psi$ | $\omega$ | **Pop (%)** |
|---|---|---|---|---|
| Cluster 1 | -79.8 (9.6) | -138.9 (13.2) | - | 100 |
| **Man $\beta$(1-4) GlcNAc** | $\phi$ | $\psi$ | $\omega$ | **Pop (%)** |
| Cluster 1 | -75.1 (17.0) | -120.5 (13.8) | - | 100 |
| **Man $\alpha$(1-3) Man (1-3)** | $\phi$ | $\psi$ | $\omega$ | **Pop (%)** |
| Cluster 1 | 72.0 (9.2) | 145.4(15.2) | - | 58.7 |
| Cluster 2 | 79.0 (12.6) | 96.5 (10.4) | - | 34.6 |
| Cluster 3 | 163.7 (11.2) | 170.9(12.0) | - | 6.7 |
| **Man $\alpha$(1-3) Man (1-6)** | $\phi$ | $\psi$ | $\omega$ | **Pop (%)** |
| Cluster 1 | 77.9 (9.4) | 156.2 (17.2) | - | 70.8 |
| Cluster 2 | 70.3 (9.4) | 108.8 (10.7) | - | 29.2 |
| **Man $\alpha$(1-6) Man** | $\phi$ | $\psi$ | $\omega$ | **Pop (%)** |
| Cluster 1 | 70.1 (10.6) | -170.6 (16.0) | 54.6 (11.1) | 82.6 |
| Cluster 2 | 83.4 (14.3) | 88.2 (12.2) | 47.8 (9.5) | 17.4 |
| **Man $\alpha$(1-6) Man (1-6)** | $\phi$ | $\psi$ | $\omega$ | **Pop (%)** |
| Cluster 1 | 70.2 (10.9) | -172.8 (18.8) | 54.8 (11.2) | 52.8 |
| Cluster 2 | 69.9 (9.7) | 176.3 (19.8) | -76.6 (12.8) | 9.5 |
| Cluster 3 | 83.1 (8.0) | -85.2 (10.9) | -80.0 (10.1) | 4.7 |
| Cluster 4 | 74.6 (15.6 | 84.7 (11.1) | 48.8 (10.8) | 16.8 |
| Cluster 5 | 69.9 (10.1) | 175.5 (21.1) | -168.1 (14.0) | 16.2 |
| **Man $\alpha$(1-2) Man (1-3)** | $\phi$ | $\psi$ | $\omega$ | **Pop (%)** |
| Cluster 1 | 76.6 (8.3) | 156.5 (13.8) | - | 77.8 |
| Cluster 2 | 68.5 (8.1) | 100.1 (9.6) | - | 22.2 |
| **Man $\alpha$(1-2) Man (1-3)** | $\phi$ | $\psi$ | $\omega$ | **Pop (%)** |
| Cluster 1 | 74.4 (9.0) | 154.8 (16.1) | - | 93.9 |
| Cluster 2 | 66.6 (4.8) | 107.4 (5.2) | - | 6.1 |
| **Man $\alpha$(1-2) Man (1-6)** | $\phi$ | $\psi$ | $\omega$ | **Pop (%)** |
| Cluster 1 | 71.8 (7.6) | 146.4 (12.9) | - | 79.8 |
| Cluster 2 | 70.0 (7.4) | 111.1 (10.1) | - | 20.2 |
| **Man $\alpha$(1-2) Man (1-6)(1-6)** | $\phi$ | $\omega$ | $\psi$ | **Pop (%)** |
| Cluster 1 | 74.7 (8.7) | 150.9 (14.9) | - | 73.8 |
| Cluster 2 | 70.4 (8.3) | 105.3 (11.6) | - | 26.2 |

### 18 DBSCAN Parameters

| Dihedral Angle's | eps ($\epsilon$) | Minimum points |
|---|---|---|
| GlcNAc $\beta$(1-4) GlcNAc | 7.5 | 75 |
| Man $\beta$(1-4) GlcNAc | 7.5 | 100 |
| Man $\alpha$(1-3) Man (1-3) | 10 | 1000 |
| Man $\alpha$(1-3) Man (1-6) | 10 | 1000 |
| Man $\alpha$(1-6) Man | 16 | 400 |
| Man $\alpha$(1-6) Man (1-6) | 16 | 400 |
| Man $\alpha$(1-2) Man | 10 | 1000 |
| Man $\alpha$(1-2) Man (2) | 10 | 1000 |
| Man $\alpha$(1-2) Man (3) | 10 | 1000 |
| Man $\alpha$(1-2) Man (4) | 10 | 1000 |

Table 17: List of parameters used for the clustering analysis of Man 9. Note: some of these parameters may need to be tweaked as clusters can bleed into each other. This algorithm will take all RAM it's given, best practice to test on smaller data sets before scaling up to production datasets in order to prevent crashes.

### 19 Man 9 / 8(II) Distance Measurements

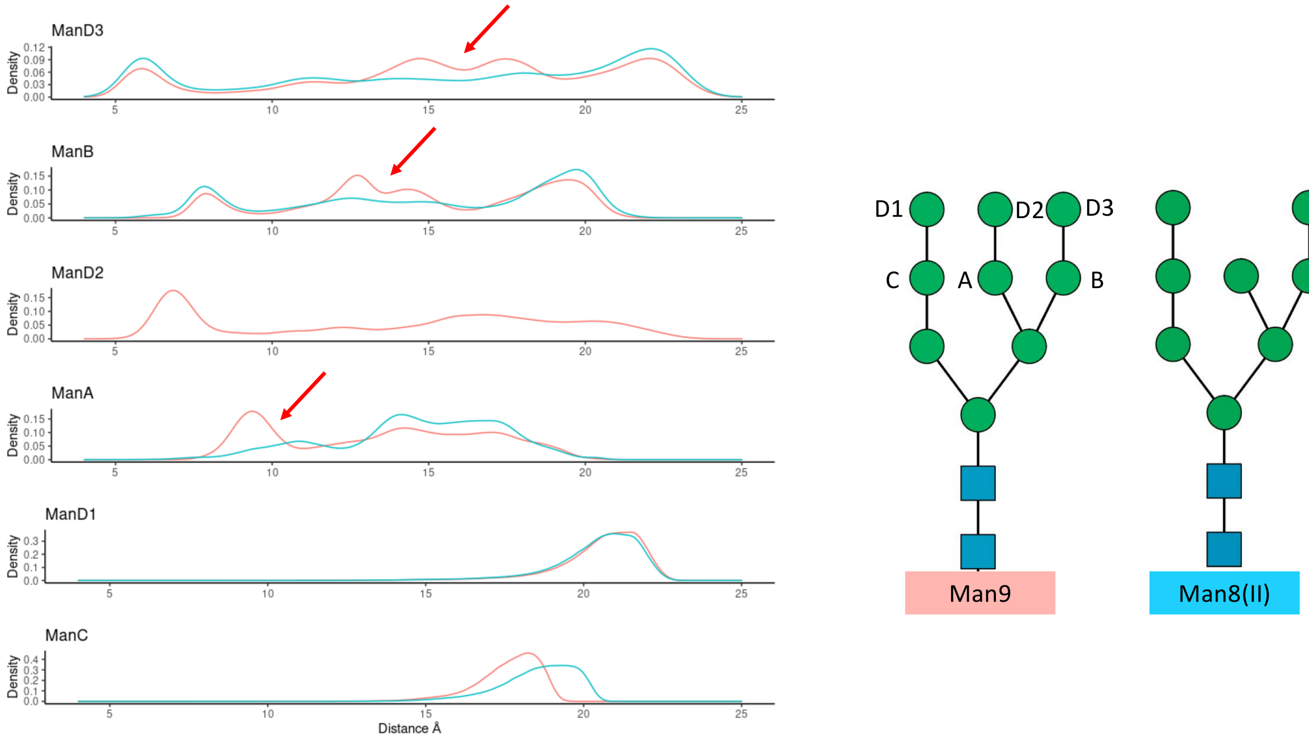


Figure 18: KDE analysis of the distances between the anomeric protons of the reducing GlcNAc and that of the specified mannose residues in the legend for Man9 (red) and Man8(II) (blue) obtained from our simulations. The red arrows highlight shorter distances only observed in Man9, which indicate the higher occurrence of folded structures, in agreement with a progressively higher occurrence of arm-arm interactions with arm elongation, as described in the main text. KDE analysis made with r and diagrams with RStudio (www.rstudio.com).
